# Pro-domain-dependent folding and co-receptor-mediated targeting to optimize an antagonistic TGF-β monomer for gene-based delivery

**DOI:** 10.64898/2026.03.23.713733

**Authors:** Łukasz Wieteska, Cynthia S. Hinck, Ananya Mukundan, Troy Krzysiak, Maarten van Dinther, Thibault Vantieghem, Rick M. Maizels, Peter ten Dijke, Caroline S. Hill, Andrew P. Hinck

**Affiliations:** Department of Structural Biology, University of Pittsburgh, Pittsburgh, PA, 15261 U.S.A.; Developmental Signalling Laboratory, The Francis Crick Institute, London NW1 1AT, U.K.; Oncode Institute and Department of Cell and Chemical Biology, Leiden University Medical Center, Leiden 2300 RC, The Netherlands; Centre for Parasitology, School of Infection and Immunity, University of Glasgow, Glasgow, G12 8TA, U.K.

**Keywords:** TGF-β, inhibition, cancer, immunosuppression, immunotherapy, tissue fibrosis

## Abstract

Transforming growth factor-beta (TGF-β), a potent promoter of extracellular matrix deposition and suppressor of infiltrating immunity, has arisen as an attractive target for improving outcomes in tissue fibrosis and cancer immune therapy. Despite the promise of TGF-β inhibitors for attenuating the progression of fibrotic disorders or as adjuncts for cancer immunotherapy, current systemically administered inhibitors that target the ligand or receptors have significant on-target liabilities, including cardiotoxicity and development of pre-malignant cutaneous squamous lesions. Recently, an engineered mini monomer of TGF-β (mmTGF-β), which potently and specifically inhibits TGF-β activity, was shown to strongly synergize with checkpoint inhibitors to suppress cancer progression in an aggressive model of melanoma when genetically delivered using an engineered form of vaccinia virus that preferentially infects cancer cells. Despite these promising results, however, a significant fraction of the mmTGF-β was found to misfold, likely due to mispairing of the cysteines that comprise its cystine knot. Here, we demonstrate that inclusion of a modified form of the TGF-β pro-domain that lacks its dimerization motif, the bowtie knot, dramatically improves both the folding and inhibitory activity upon secretion by mammalian cells, thus overcoming one of the major limitations of genetically delivering mmTGF-β. Furthermore, we show that fusion of mmTGF-β to a CD44 binding domain enhances the inhibitory potential of mmTGF-β on immune cells, and on other cell types which express CD44, by more than 30-fold compared to cells negative for CD44. Together, these modifications provide a framework for further enhancing the efficacy and safety of mmTGF-β for cancer immune therapy, and possibly also tissue fibrosis, when delivered genetically using vaccinia, or other related approaches.

## Introduction

Transforming growth factor-beta 1, 2, and 3 (TGF-β1, -β2, and –β3) are secreted signalling proteins comprised of two cystine-knotted growth factor domains joined together by a single inter-chain disulfide bond^1^. They play essential roles in tissue maintenance, where they potently stimulate the expression of extracellular matrix (ECM) proteins, such as collagen and fibronectin^2^, and epithelial-to-mesenchymal cell transition (EMT)^3^, which is required for tissue remodeling and maintenance of vital tissues, such as the heart^4–7^. TGF-bs also suppress tumor growth by inhibiting epithelial cell proliferation^8,9^, but promote immune tolerance by upregulating suppressive regulatory T-cells (T_regs_) and downregulating B-cells^10,11^.

The TGFβs are synthesized as pro-cytokines^12^, and only after processing by pro-protein convertases during secretion and release from their inhibitory prodomain^13–15^, are able to bind and assemble their signalling receptors, two structurally similar, but functionally distinct, single-pass transmembrane receptor kinases, into a signalling-competent (TGFBR1)_2_(TGFBR2)_2_ heterotetramer^16,17^. The TGFβs assemble their signalling receptors in a stepwise manner, first by binding TGFBR2 and then by binding and recruiting TGFBR1, which is due to TGFBR1 binding to a composite interface formed by both monomers of TGF-β and the TGFBR2 extracellular domain (ECD)^13,18^.

In humans, dysregulation of TGF-β signalling can promote the progression of fibrotic disorders, such as idiopathic pulmonary fibrosis (IPF), renal fibrosis, and cardiac fibrosis^19^, but also most soft-tissue cancers^9 10^. In the former setting, tissue inflammation and other processes can increase the release of bioactive TGF-β, which can drive disease progression through the expression and excessive deposition of the extracellular matrix proteins. In the latter setting, cancer cells can lose sensitivity to TGF-β growth inhibition by dysregulation of the cell cycle or loss of essential pathway components, such as SMAD4, while the surrounding stromal cells, including cancer associated fibroblasts (CAFs) and immune cells do not. This can enable tumor immunity and can contribute to immune exclusion in tumors and the tumor microenvironment (TME)^20,21^, a major barrier for the success of immune checkpoint, chimeric antigen receptor T-cell (CAR-T), and other immune therapies, which even with their limitations, have revolutionized the treatment of cancer^22–24^. TGF-β has therefore emerged as an important target for fibrotic disorders and cancer and different classes of inhibitors that target the pathway are being aggressively pursued in clinical studies, as monotherapies and as adjuncts to other therapies, such as cancer immune therapy^25,26^.

One of the major challenges associated with inhibiting TGF-β stems from its pleiotropy, as it plays essential roles in tumor suppression, tolerogenic immune signalling, and tissue homeostasis in a variety of organ systems^8,9^. Moreover, a considerable quantity of TGF-β exists in a latent form in the extracellular matrix (ECM) bound to its pro-domain and to latent TGF-β binding proteins (LTBPs)^27^. Attempts to neutralize mature TGF-β using monoclonal antibodies or receptor traps, or to block the receptors with antibodies or small molecule kinase inhibitors, have resulted in considerable dose-limiting toxicities, including on-target cardiotoxicity, development of keratocanthomas, pre-malignant cutaneous squamous lesions, and persistent haemorrhagic bleeding ^6,7,28–31^.

Recently, the underlying structural basis by which TGF-β engages its receptors and assembles them into an active signalling complex in a stepwise manner^18,32^ has been leveraged to develop an engineered mini monomer of TGF-β (hereafter, mmTGF-β) that binds TGFBR2, but is unable to bind and recruit TGFBR1^33^. This highly selective and potent inhibitor can compete with native TGF-β for binding to cell surface TGFBR2, thereby inhibiting TGF-β1, -β2, and –β3 signalling without targeting TGF-β itself. In our published data, we have shown that recombinant mmTGF-β delivered with osmotic pumps in an animal model of carcinogen-induced oral cancer is highly effective in attenuating disease progression^34^. In addition, we have also taken advantage of the genetically encoded nature of mmTGF-β to deliver it to tumors and the TME using oncolytic viruses (OVs) engineered to express mmTGF-β and have shown that this has superior therapeutic effects compared to checkpoint inhibitors alone in animal models of checkpoint refractory melanoma^35^. Importantly mmTGF-β has some very unique properties, including its small size (ca. 10 kDa) and its specific targeting of TGFBR2 that might allow it to overcome some of the shortcomings of other classes of inhibitors, such as neutralizing antibodies, receptor traps, or TGFBR1 kinase inhibitors.

Owing to TGF-β pleiotropy and the potential for side effects arising from on-target inhibition in non-target tissues, but also challenges that would be faced delivering recombinant mmTGF-β, such as difficulty producing large quantities of the highly disulfide-bonded mmTGF-β for therapeutic use and the likely requirement to fuse to a fragment crystallizable (F_c_) domain or albumin to increase its circulation half-life, we wanted to build upon our prior success in delivering mmTGF-β genetically. Previously, we showed that infection of tumor cells by recombinant OVs bearing mmTGF-β results in the release of active, inhibitory mmTGF-β, although much of the secreted protein appears to be non-native and non-inhibitory as a result of the mmTGF-β forming disulfide-linked aggregates when expressed in mammalian cells. Here, we aimed to improve the folding and activity of mmTGF-β, by producing mmTGF-β constructs with either modification or simplification of its disulfide structure, or inclusion of an engineered form of the TGF-β pro-domain. We found that while the former can confer modest improvements in folding, much more significant improvement can be achieved by inclusion of an engineered pro-domain. We have also fused mmTGF-β with a CD44 co-receptor extracellular domain binding module and found that this greatly increased potency against target cells expressing CD44 compared to cells that are null for CD44. This has the potential to increase safety by biasing the inhibitory effect to different classes of immune cells that abundantly express the co-receptor CD44. Together, these modifications promise to expand the efficacy and safety of mmTGF-β as a therapeutic agent when delivered genetically.

## Results

### mmTGF-β design and improvement of its solubility

mmTGF-β is derived from the mature TGF-β2 ligand and contains modifications that eliminate the cysteine residue (C77S) responsible for forming the native disulfide-linked TGF-β homodimer and replace the heel helix (Δ52–71) with a flexible loop (**Figure S1A**). These modifications convert the native homodimeric TGF-β2 agonist, which normally engages two type I and two type II receptors, TGFBR1 and TGFBR2, respectively, into a monomeric ligand^33^. The consequence of these modifications is that mmTGF-β selectively binds TGFBR2, but cannot bind and recruit TGFBR1, thereby functioning as a competitive inhibitor of TGF-β signalling. To impart high-affinity binding to TGFBR2 comparable to that of TGF-β1 and TGF-β3, and thus effective antagonism against native TGF-β dimers, the original mmTGF-β2 construct incorporated seven additional substitutions, K25R, R26K, I89V, I92V, N94R, T95K, and I98V, so that the residues in the binding interface with TGFBR2 matched those of TGF-β1 and TGF-β3, which have a higher affinity for TGFBR2 than does TGF-β2^33^.

In its initial conception, the modified mmTGF-β inhibitor, known as mmTGF-β2-7M, was found to exhibit greatly improved solubility compared to native dimeric TGF-β1, -β2, and – β3, which are only soluble at micromolar or lower concentrations at neutral pH^36^. To further improve the solubility of mmTGF-β2-7M as an inhibitor, which we believe is important for effective antagonism, we introduced two additional substitutions in the loop replacing the heel helix: S57R and S59R (2R) (**Figure S1A**). These substitutions resulted in a modest increase in signal amplitude and reduced peak broadening in the NMR spectra at both pH 4.6 and 7.2 (**Figure S1B, C**), suggesting a reduced propensity to form soluble aggregates. In light of the improved hydrodynamic behavior of the new variant with S57R and S59R, which we designate as mmTGF-β2-7M-2R, we have used this as the basis for further development as described below.

### Inefficient folding of mmTGF-β compromises its recombinant production

mmTGF-β2-7M-2R (hereafter mmTGF-β), was successfully employed in a mouse oral squamous cell carcinoma model using sustained systemic delivery via subcutaneous administration^34^, and subsequently in a head and neck and melanoma mouse model^35^. In both settings, mmTGF-β synergized with anti-PD-L1 and anti-PD-1 immunotherapy and demonstrated promising therapeutic efficacy. In the first instance, mmTGF-β was produced in bacteria, refolded from inclusion bodies, and purified to homogeneity, whereas in the second, mmTGF-β was produced by the host mouse cells targeted by the recombinant vaccinia virus. Despite clear functional efficacy when mmTGF-β was produced *in situ* by tumor cells following vaccinia virus delivery^35^, Western blot analysis under non-reducing conditions revealed that recombinant expression in HEK293 cells predominantly led to the secretion of disulfide-linked aggregates, whereas the monomeric form, which may correspond to the active form of the protein, was barely detectable or not detectable (**Figure S2A**). This prompted us to refine the design of this inhibitor to improve its folding and facilitate its deployment as a therapeutic via gene-encoded delivery. In parallel, we sought to evaluate the feasibility of incorporating the inhibitor into multidomain architectures to achieve cell-type–specific targeting.

### Disulfide bond simplification reveals essential and non-essential bonds in mmTGF-β

Growth factors of the TGF-β family contain three conserved intramolecular disulfide bonds that form a cystine knot core^37^ (**Figure 1A**). In addition, some TGF-β family members possess an extra intrachain disulfide bond, referred to here as the auxiliary disulfide (A), which tethers the N-terminal α1-helix to the remainder of the cystine knot domain (**Figure 1A**). Because mmTGF-β expressed in mammalian cells appeared to accumulate as disulfide-linked aggregates, we hypothesised that reducing the number of disulfide bonds might alleviate misfolding. To test this, individual disulfide bonds were systematically removed by cysteine-to-alanine substitutions, and the folding and activity of each variant were assessed by NMR and functional assays. Removal of the auxiliary disulfide (mmTGF-β-ΔA) resulted in a protein that could be folded efficiently, exhibited a well-dispersed ^1^H-^15^N NMR spectrum consistent with a native-like structure (**Figure 1B**), and retained inhibitory activity in a HEK293T cell-based CAGA_12_-luciferase reporter assay, which is sensitive to the induction of pSMAD3–SMAD4 complexes by TGF-β^38^ (**Figure 1C**). In contrast, replacement of any of the three disulfide bonds forming the cystine knot core led to misfolding and loss of function (**Figure 1B**). Removal of the auxiliary disulfide resulted in a slight decrease in thermal stability, as reflected by differential scanning calorimetry (**Figure S2B**). However, due to the intrinsically high stability of the disulfide core, the measured melting temperature remained high (∼76.5°C for mmTGF-β-ΔA, compared to ∼81.8°C for mmTGF-β).

**Figure 1.**
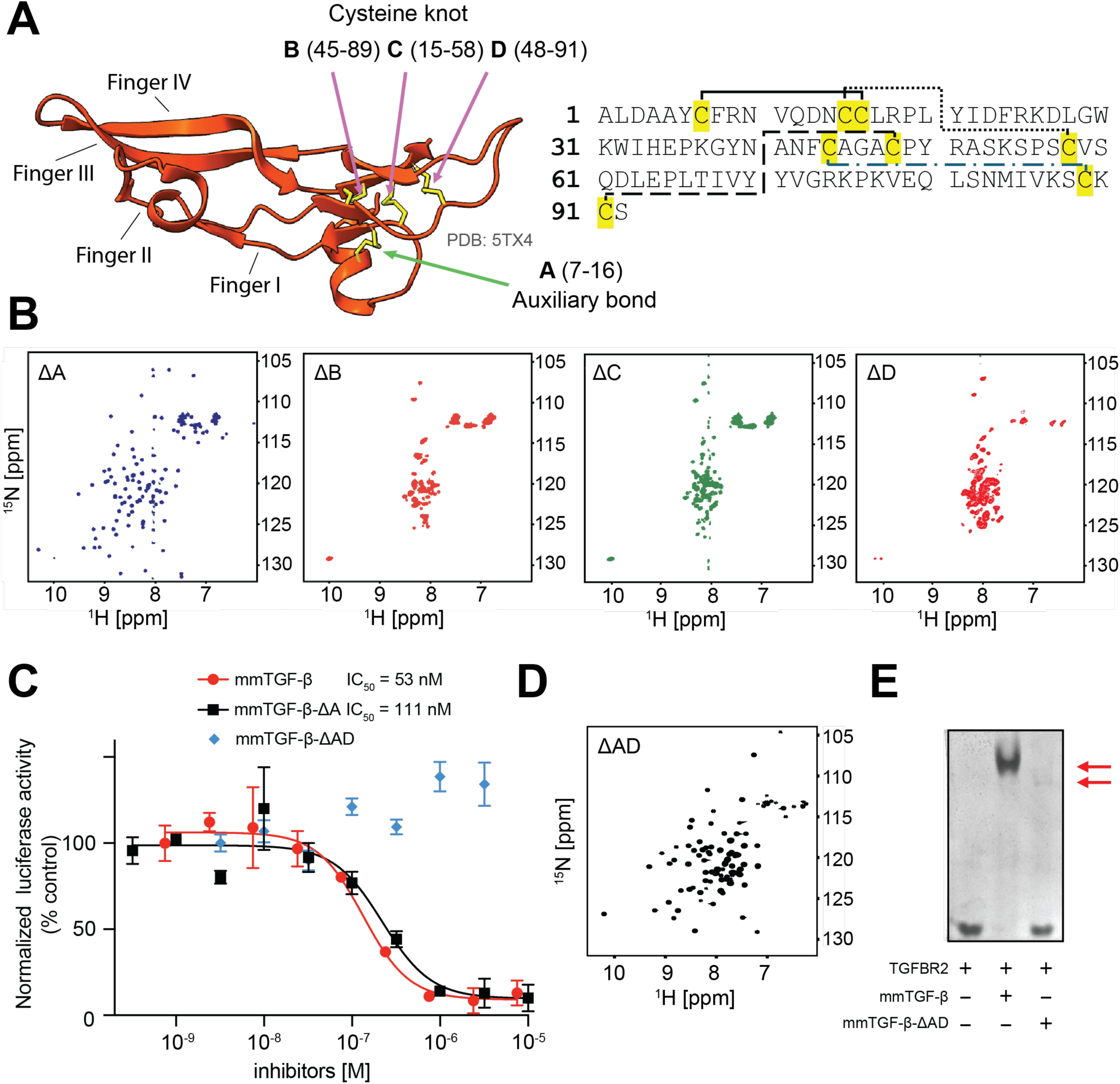
Simplification of the disulfide core. A. Spatial arrangement of cysteine residues in mmTGF-β illustrated using the structure of the original inhibitor (PDB: 5TX4), alongside their connectivity and positions within the amino-acid sequence. B. The amide region of the ¹H-¹⁵N correlation spectra showing the eNects of individual cysteine-to-alanine substitutions in mmTGF-β. Only disruption of the auxiliary disulfide bond (“A”) yielded a folded protein (mmTGF-β-DA), whereas substitutions within the cystine knot prevented proper folding. C. Inhibitory activity of mmTGF-β, mmTGF-β-ΔA and mmTGF-β-ΔAD, measured using HEK293 cells stably expressing the CAGA_12_-luciferase/Renilla reporter. Inhibitors were tested at increasing concentrations in the presence of a constant TGF-β1 concentration and increasing inhibitor doses. Both mmTGF-β and mmTGF-β-ΔA show comparable activity, whereas mmTGF-β-ΔAD showed no detectable activity. Presented curves are fitted to datapoints collected from two independent repeats with three technical replicates each. D. The amide region of the ¹H-¹⁵N correlation spectra showing the successful substitution of bonds “A” and “D” in mmTGF-β. Spectra were recorded with the addition of 1mM CHAPS E. Native PAGE analysis of complex formation between mmTGF-β, mmTGF-β-ΔAD, and the extracellular domain of TGFBR2. Upper bands correspond to protein–protein complexes (red arrows), which is prominent for mmTGF-β but markedly reduced for mmTGF-β-ΔAD, suggesting significantly weaker binding.

To explore whether alternative stabilizing interactions could partially compensate for the loss of a cystine knot disulfide, we focused on disulfide bond “D”, which is positioned furthest from the receptor-binding finger regions and has a redundant structural role with bond “C” in tethering the C-terminal strand to one of the β-strands that forms finger 3. Several residue-pair substitutions were screened using yeast surface display. Among the tested variants, one construct showed improved expression and folding on the yeast cell surface (**Figure S2C**), prompting further characterization. Interestingly, this variant (mmTGF-β-ΔAD) could be refolded into a native TGF-β-like structure, as evidenced by ^1^H-^15^N NMR (**Figure 1D**). However, despite adopting a folded conformation, this variant displayed impaired binding to TGFBR2, as assessed by native gel electrophoresis (**Figure 1E**), and was inactive in the CAGA_12_-luciferase SMAD3 transcriptional reporter assay (**Figure 1C**).

Together, these results demonstrate that while the auxiliary disulfide bond can be removed without compromising folding or activity, the integrity of the three disulfides that comprise the cystine knot is essential not only for structural stability, but also for productive receptor engagement.

### Use of the PRDC-based cysteine knot scaffold

As an alternative strategy to overcome folding limitations, we reasoned that because some cystine-knot growth factors, like sclerostin and PRDC, are naturally produced without pro-domains, their cores may be intrinsically more permissive to folding^39^. Building on this observation, we designed two PRDC-based chimeras: one in which the mmTGF-β fingers were grafted onto the full PRDC scaffold (mmTGF-β-PRDC), and a second, in which only the PRDC cystine knot core was retained (mmTGF-β-PRDCs) (**Figure 2A**). Both constructs were produced in *E. coli* as inclusion bodies, refolded, and purified to homogeneity. Samples of both constructs exhibited well-dispersed ^1^H-^15^N NMR spectra indicative of natively folded proteins (**Figure 2B**). The longer construct showed several unresolved peaks in the center of the spectrum, most likely attributable to an unstructured N-terminal extension and a long unstructured internal loop (**Figure 2A**). Both constructs displayed robust inhibitory activity in HEK293 cells, although mmTGF-β-PRDC showed slightly reduced potency (**Figure 2C**).

**Figure 2.**
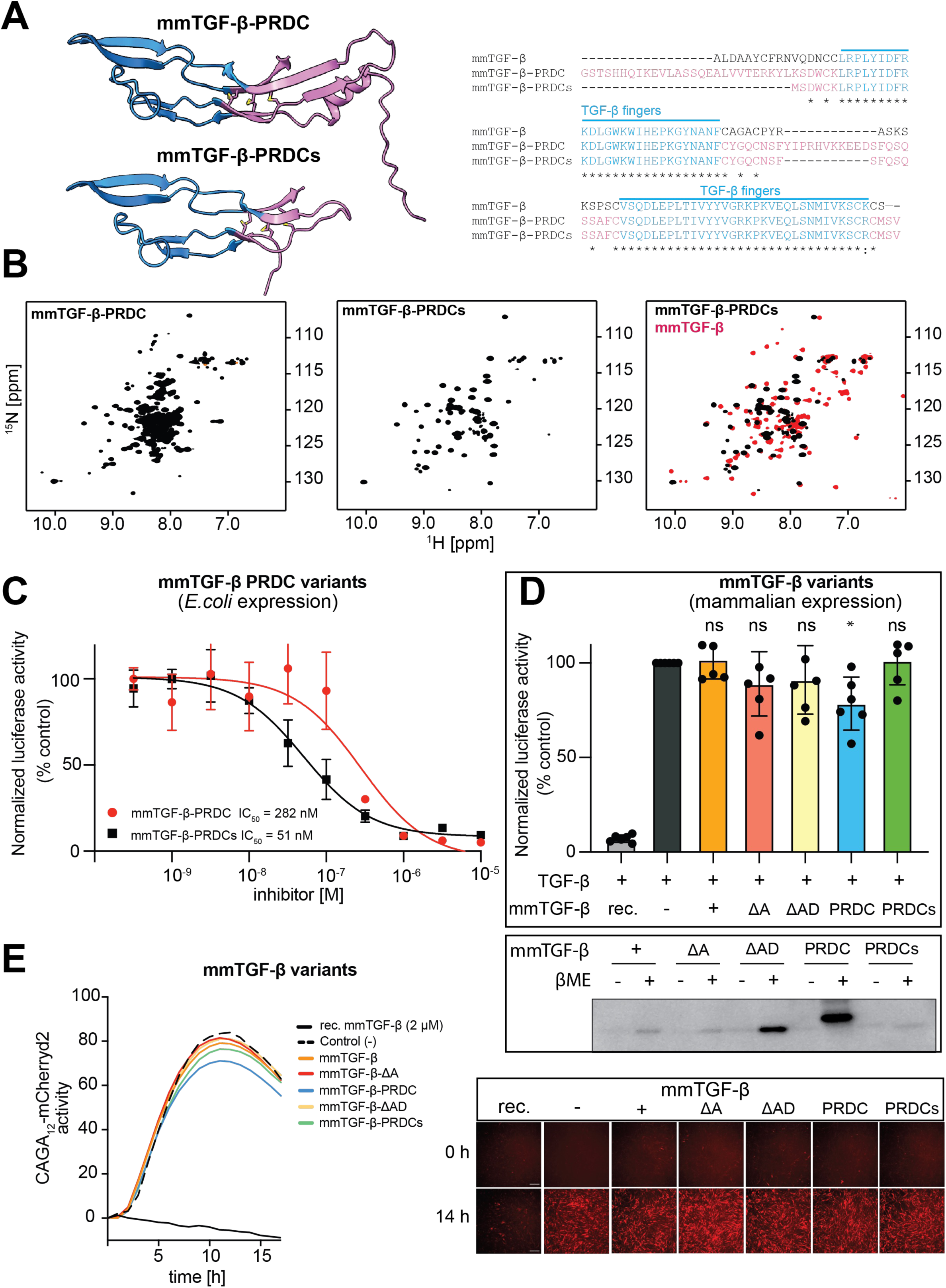
Incorporation of alternative core sca;olds from PRDC, a growth factor devoid of a prodomain. A. AlphaFold3 models of mmTGF-β-PRDC and its shorter variant mmTGF-β-PRDCs, with the corresponding sequence alignment shown on the right. Parallel β-strands forming the TGF-β fingers are depicted in blue, while the cystine core with N terminal portion grafted from human PRDC is highlighted in pink. Asterisk indicate identity. B. The amide region of the ¹H-¹⁵N correlation spectra for mmTGF-β-PRDC and mmTGF-β-PRDCs. Both spectra indicate that the proteins are folded. In mmTGF-β-PRDC, a cluster of unresolved peaks between 8 and 9 ppm in the ¹H dimension suggests the presence of a coiled-coil like region, likely arising from the extended N-terminal segment. The right-most panel shows an overlay of mmTGF-β-PRDC (black) with mmTGF-β (red). The close correspondence of many peak positions supports a largely conserved overall fold. C. Inhibitory activity of mmTGF-β-PRDC and mmTGF-β-PRDCs, measured using the dual CAGA_12_-luciferase/Renilla reporter assay at a constant TGF-β1 concentration and increasing inhibitor doses. Both inhibitors demonstrate activity. The presented curves are fitted to datapoints collected from two independent repeats with three technical replicates each. D. Inhibitory activity of conditioned media containing secreted mmTGF-β variants in the MDA-MB-231 CAGA_12_-luciferase/Renilla reporter assay. Conditioned media from HEK293 cells transiently transfected with consecutive variants were analysed at a single 16h endpoint. The experiment was performed at least three times; the data shown are derived from two biological repeats, each with three technical replicates. Statistical analysis was performed using GraphPad Prism (version 10.4.1). Data were analysed using one-way ANOVA. The bottom panel shows Western blot analysis of mmTGF-β variant expression (anti-HA tag). mmTGF-β-ΔAD and mmTGF-β-PRDCs exhibited the highest expression levels; however, bands were predominantly detected under reducing conditions, indicating impaired folding during mammalian expression. E. Inhibitory activity assessed in the NIH3T3 CAGA_12_-mCHERRYd2 reporter line, monitored hourly for 16h following TGF-β1 addition. Representative fluorescent micrographs of NIH3T3 cells expressing mCHERRYd2 upon 2 ng/ml TGF-β1 stimulation are shown below. The experiment was repeated at least three times. The presented data were analysed from two biological repeats, each comprising six technical replicates and two ROIs per replicate.

### Assessment of construct expression and activity in mammalian expression systems

Next, to determine which design strategy yielded improved production and folding in mammalian cells, all constructs were re-cloned into the mammalian expression vector pcDNA3.1+ with an N-terminal hemagglutinin (HA) tag to enable detection and expressed in suspension-cultured HEK293 (expi293) cells. Protein expression and inhibitory activity in the conditioned media were assessed by Western blotting and functional assays, respectively, with the activity measured in two reporter cell lines: MDA-MB-231 cells stably expressing the CAGA_12_-luciferase reporter and NIH3T3 cells expressing CAGA_12_-mCHERRYd2 reporter. As shown, most of the tested conditioned media did not exhibit significant inhibitory activity towards the TGF-β1 stimuli (**Figure 2D, E)**. This is consistent with the detection of low levels of monomers in Western blots under non-reducing conditions (**Figure 2D**), with the majority of the protein becoming detectable only after reduction with β-mercaptoethanol, suggesting inefficient folding. However, one of the constructs, mmTGF-β-PRDC, stood out from the rest, exhibiting a higher expression level than the other variants and a low level of inhibitory activity, which was statistically significant (p=0.025). In this construct, the extended N-terminal region derived from the PRDC construct (17 additional residues relative to the original construct) may help to promote native folding or, alternatively, enhance expression.

### Incorporation of an engineered pro-domain enables production of an active inhibitor in mammalian cells

Strategies described above yielded at best marginal improvement in the production of native inhibitory mmTGF-β in mammalian cells, and therefore, an alternative approach was investigated. The N-terminal extension in mmTGF-β-PRDC modestly improved both protein expression and activity and appeared to function in a pro-domain–like manner. This mirrors to some extent the well-established role of the TGF-β pro-domain as an intramolecular chaperone that assists in the folding of the growth factor during biosynthesis^27^. We therefore attempted to incorporate the TGF-β pro-domain into mmTGF-β. However, the initial construct in which the full pro-domain was fused to mmTGF-β failed to yield detectable protein expression in HEK293 cells. Previous studies had demonstrated that substitution of the two bowtie cysteines with serines does not impair expression and secretion of TGF-β^40^, and thus, we designed a more restricted construct based on pro-TGF-β3. In this design, residues 227–253, responsible for forming the bowtie element were deleted (**Figure 3A**). In addition, the mmTGF-β sequence was redesigned to correspond directly to the TGF-β3 ligand sequence, thereby matching the pro-domain with the growth factor with which it naturally pairs (**Figure S3**); this new form of mmTGF-β is designated as mmTGF-β3. We generated two variants: one furin-resistant, p-mmTGF-β3-FR, in which the native furin cleavage site was substituted with a furin-resistant sequence and another, p-mmTGF-β3-2F, in which the endogenous furin site was retained and an additional furin site was introduced in a loop of the pro-domain (residues 86–89) to facilitate internal cleavage of the pro-domain and efficient inhibitor release. As shown by the anti-HA-tag Western blot, both constructs were successfully expressed (**Figure 3B**). Treatment with PNGase F, which removes N-linked glycosylation, resulted in a clear increase in band mobility, consistent with the removal of glycans attached to the three asparagine residues in the prodomain (**Figure S3**). For the cleavable construct, three bands are observed: A*, corresponding to the uncleaved p-mm-TGF-β3 (∼42 kDa); B*, corresponding to the isolated prodomain following cleavage and release of the active mmTGF-β (∼31.5 kDa); and C*, corresponding to the prodomain processed at the secondary cleavage site (∼10 kDa). The addition of a reducing agent to the furin-containing construct did not substantially increase band intensity, suggesting that the majority of the protein was correctly folded and already efficiently processed. An additional upper band, observed with the furin-resistant construct at approximately twice the molecular weight, likely represents a dimer. This species most probably arises from an interaction in which an active monomer engages with the pro-domain of a second protomer. This artifact is observed only for the furin cleavage-site construct and occurs mostly under high-expression conditions in expi293 cells (see **Figure 5B**).

**Figure 3.**
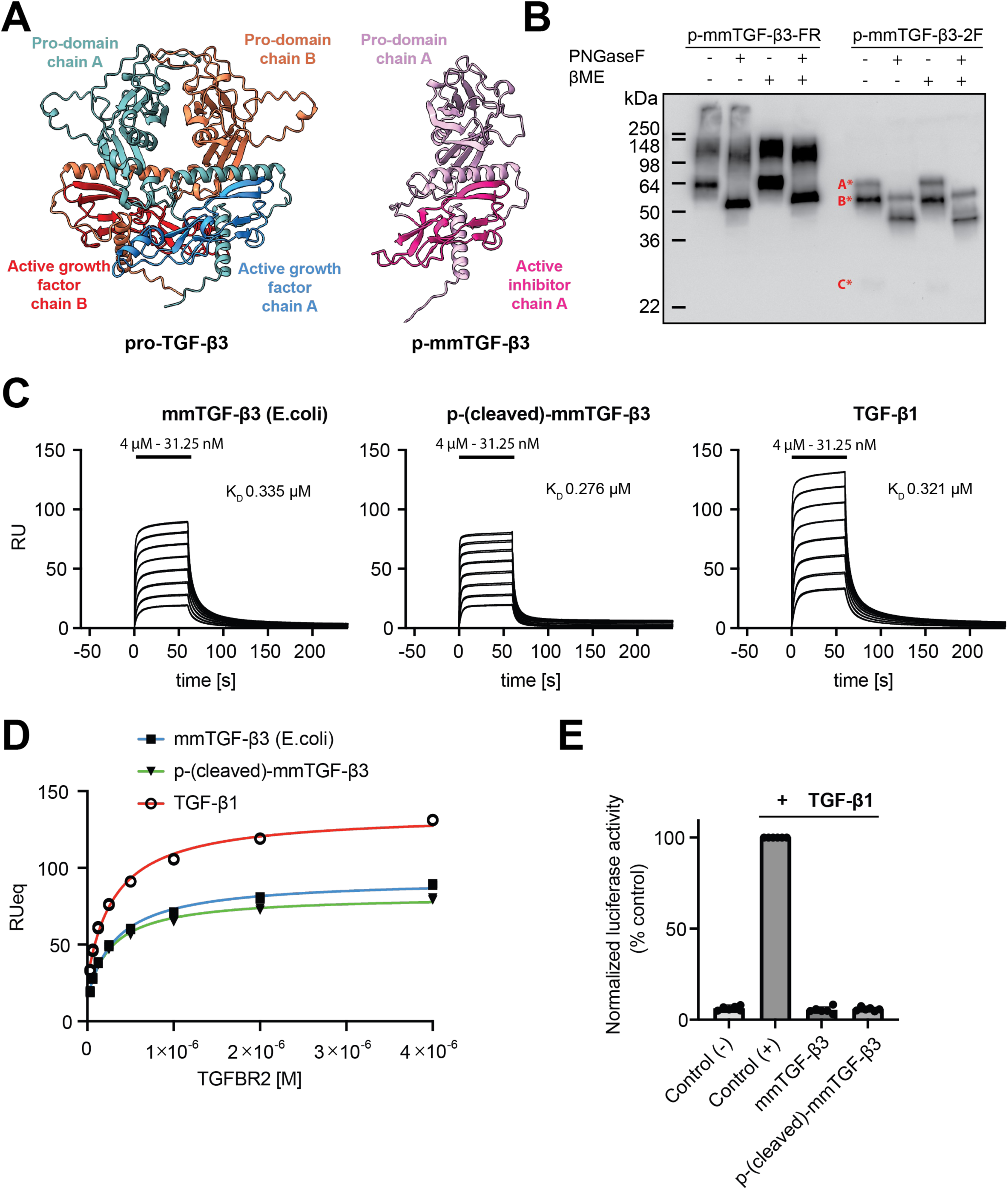
Adaptation of pro-domain fragments. A. AlphaFold3 models of pro-TGF-β3 and p-mmTGF-β, highlighting key dimeric structural features and showing how the prodomain was incorporated into the p-mmTGF-β design. B. Western blot analysis of conditioned media from HEK293 cells transfected with the furin-resistant (p-mmTGF-β-FR) and furin-cleavable (p-mmTGF-β-2F) constructs. Red-labeled dots mark the corresponding bands discussed in the main text. C, D. SPR binding sensorgrams of the extracellular domain (EC) of TGFBR2 to immobilized mmTGF-β3 (produced in *E.coli*), mmTGF-β3 active monomer derived from p-HRV3C-mmTGF-β3 and TGF-β1 on the chip surface. Sensorgrams show a 2-fold dilution series of EC TGFBR2 injections in the ranges shown (black). Fitting curves (D) are derived from data collected in three consecutive repeats. E. Inhibitory activity of mmTGF-β3 active monomer derived from p-HRV3C-mmTGF-β3 using the MDA MB-231 cell line harbouring CAGA_12_-luciferase/Renilla reporter assay, measured at a single 16 h endpoint following TGF-β1 stimulation. Both bacterially and mammalian-cell–produced mmTGF-β were tested at an identical concentration (2 µM). The presented data were analysed from two biological repeats, each comprising three technical replicates.

Next, we sought to determine whether mmTGF-β3 released from the pro-domain is indeed natively folded and inhibitory. To investigate this, we engineered a variant in which the primary furin cleavage site was replaced with a human rhinovirus 3C (HRV3C) protease cleavage site (p-mmTGF-β3-HRV3C) and the second furin site was restored to native sequence. The p-mmTGF-β3-HRV3C protein was purified from conditioned medium by IMAC chromatography, subjected to HRV3C protease cleavage, and further purified (**Figure S4A-B**). Measurement of the intact mass confirmed the expected molecular weight and formation of all disulfide bonds. Functional activity of the purified mmTGF-β3 was demonstrated by binding to recombinant TGFBR2 ectodomain using SPR (**Figure 3C–D**) and by inhibition of TGF-β-stimulated CAGA_12_-luciferase reporter in MDA-MB-231 cells (**Figure 3E**). Additionally, we confirmed that mmTGF-β3, similarly to mmTGF-β, can be refolded from bacterially produced inclusion bodies and possesses potent inhibitory activity in a CAGA_12_-luciferase reporter assay in MDA-MB231 cells (**Figure S4 C-D**).

Next, we evaluated the activity of p-mmTGF-β3-2F in HaCaT cells stably expressing a GFP-SMAD2 reporter^41^ (**Figure 4A–C**). Upon stimulation with TGF-β1, SMAD2 is phosphorylated by the activated TGFBR1 kinase, forms homo- or heterotrimeric complexes with SMAD4, and rapidly accumulates in the nucleus^42^. This process can be readily observed and quantified in live cells. Pre-treatment with 2 µM mmTGF-β completely abolished TGF-β1 induced signalling, as evidenced by the absence of GFP-SMAD2 nuclear accumulation. A comparable inhibitory effect was observed when cells were treated with conditioned medium containing p-mmTGF-β3-2F. There was no effect on TGF-β-induced GFP-SMAD2 nuclear accumulation when conditioned medium containing the furin-resistant variant, p-mmTGF-β3-FR was used or with the conditioned medium alone (Control). These findings were independently confirmed using NIH3T3 cells harboring a CAGA_12_-mCHERRYd2 reporter (**Figure 4D**).

**Figure 4.**
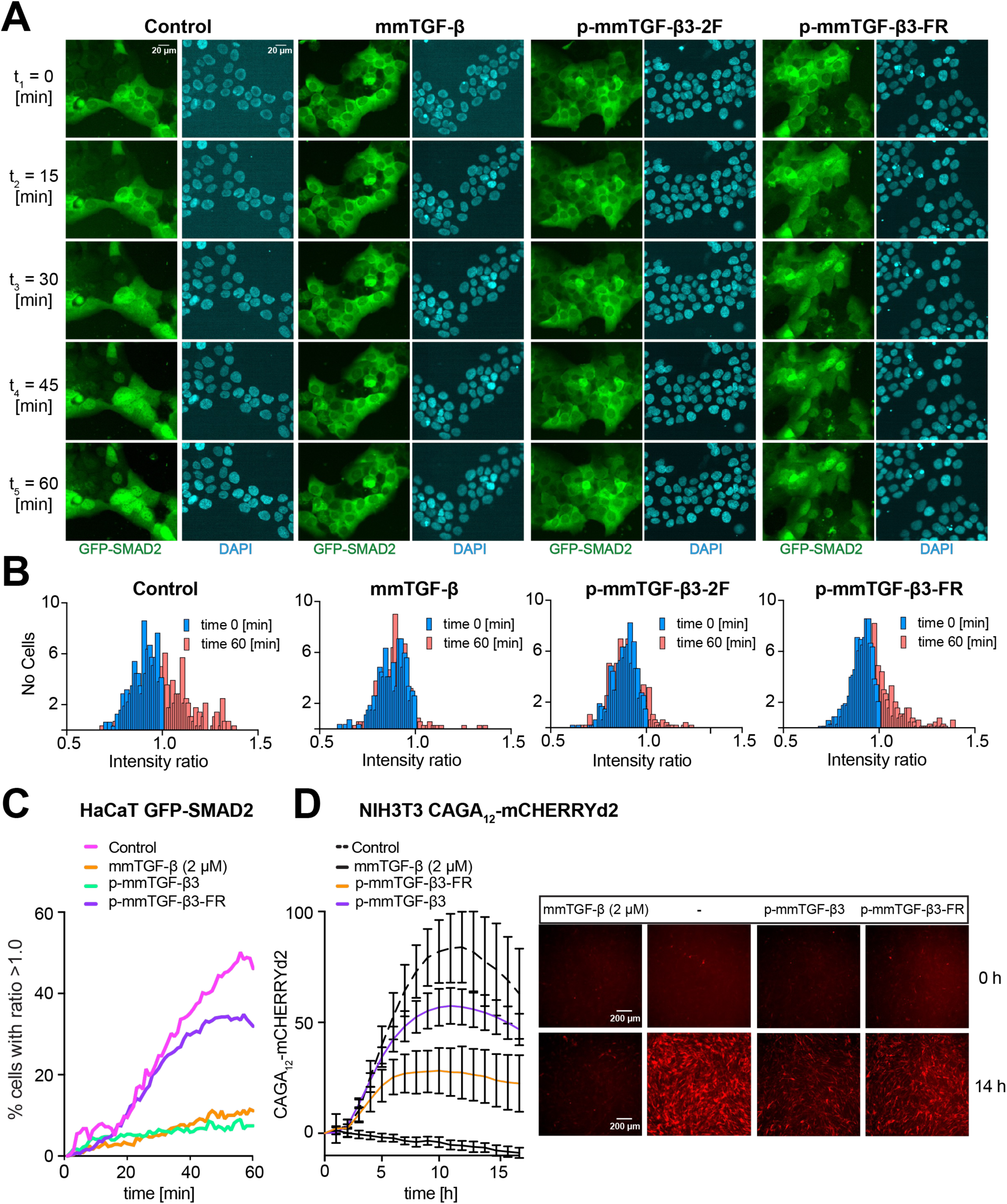
Active mmTGF-β is released from its pro-domain and actively inhibits TGF-β signalling. A–C. GFP-SMAD2 nuclear accumulation assay was used to assess inhibition of the TGF-β pathway. Conditioned media containing p-mmTGF-β or p-mmTGF-β-FR (furin-resistant) were tested alongside bacterially produced mmTGF-β (positive control). As shown in the micrographs (A) and accompanying analyses (B–C), cells treated with either conditioned media or mmTGF-β did not exhibit GFP–SMAD2 nuclear accumulation upon TGF-β1 (2 ng/ml) stimulation. Quantification at the 60-minute endpoint (B) shows the proportion of cells displaying altered nuclear-to-cytoplasmic GFP–SMAD2 ratios, reflecting inhibition of SMAD2 nuclear accumulation. The time-course analysis (C) reveals that divergence from the control becomes detectable after ∼20 minutes and reaches its maximum approximately 30 minutes later. The experiment was performed several times. The presented data were analysed from two biological repeats, each comprising four technical replicates. D. Inhibitory activity of conditioned media containing secreted p-mmTGF-β and p-mmTGF-FR variants. Conditioned media from HEK293 cultures transiently transfected with consecutive variants were tested using the NIH3T3 CAGA_12_–mCHERRYd2 reporter line, and the cells were monitored hourly for 16 h following addition of TGF-β. On the bottom panel selected fluorescent micrographs of NIH3T3 cells expressing mCHERRYd2 upon TGF-β1 stimulation (2 ng/ml).

### A single furin cleavage site is sufficient for the release of active mmTGF-β3 inhibitor

Physiologically, proteolytic processing at the furin site is not sufficient to generate freely active TGF-β ligand, because the mature growth factor remains tightly, non-covalently associated with the latency-associated peptide (LAP), which occludes receptor-binding surfaces and maintains the ligand in a completely latent state^43^. *In vivo*, productive activation typically requires additional mechanisms that disrupt or remodel the LAP “cage”, such as integrin-mediated mechanical force^14,44^ or protease-dependent pathways^45^ that deform the latent complex or cleave the pro-domain to expose or release the mature growth factor. Given this, we were concerned that a single furin site might not enable release of active mmTGF-β3 from the engineered pro-domain due to pro-domain:mmTGF-β3 binding. We therefore generated additional constructs containing zero, one, or three furin sites to complement the existing construct with two furin sites (**Figure 5A**). These constructs were each successfully expressed (**Figure 5C**) and apart from mmTGF-β3-FR, all variants displayed inhibitory activity in the CAGA_12_-luciferase assay in MDA-MB-231 cells (**Figure 5B**). Though the differences between the –1F, -2F, and –3F constructs were not statistically significant, the highest apparent potency was observed for mmTGF-β3-1F, suggesting that processing at engineered sites in addition to the single native furin site separating the pro-and GF domains is not required to enable the full inhibitory capacity of mmTGF-β3.

**Figure 5.**
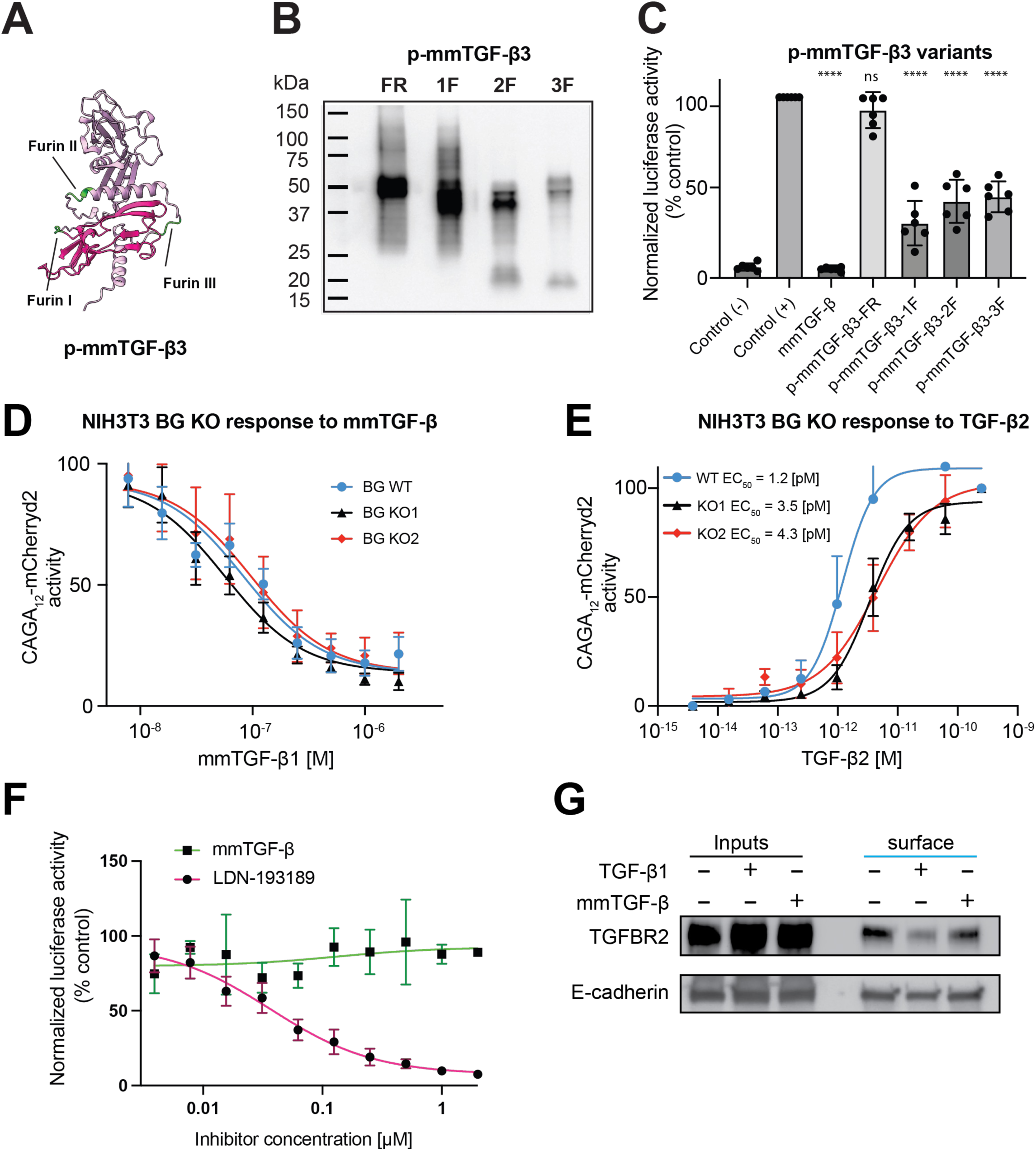
A single furin cleavage site is su;icient for p-mmTGF activity, that is TGF-β pathway specific, co-receptor independent and does not trigger receptor internalisation. A. AlphaFold3 model of p-mmTGF-β illustrating the locations of the engineered furin cleavage sites within the construct B. Western blot analysis of mammalian expression of p-mmTGF-β variants engineered to contain between zero and three furin cleavage sites (FR, 1F,2F and 3F). C. Inhibitory activity of conditioned media containing of secreted p-mmTGF-β variants (B). Conditioned media from HEK293 cultures transiently transfected with consecutive variants were tested using the MDA MB-231 cell line with CAGA_12_-luciferase/Renilla reporter assay, measured at a single 16 h endpoint following TGF-β1 stimulation. The presented data were analysed from three biological repeats, each comprising two technical replicates. Statistical analysis was performed using GraphPad Prism (version 10.4.1). Data were analysed using one-way ANOVA. D, E. The inhibitory activity of mmTGF-β was assessed in wild-type NIH3T3 cells and two betaglycan-knockout (BG KO) clones carrying the CAGA_12_-mCHERRYd2 reporter. mmTGF-β showed comparable inhibitory potency in wild-type and BG KO cells, demonstrating that its activity is independent of betaglycan (A, C). In contrast, cellular responses to low concentrations of TGF-β2 are potentiated by betaglycan (B), as expected. E. Selectivity toward TGF-β signalling was assessed by treating MDA-MB-231 cells harbouring a BRE-luciferase/Renilla reporter with increasing concentrations of mmTGF-β or the BMP type I receptor kinase inhibitor LDN-193189. BMP4 stimulation across the tested concentration range of mmTGF-β was not impacted, whereas LDN-193189 produced dose-dependent inhibition, with an IC₅₀ of 37 ±14 nM, consistent with the literature. F. A surface biotinylation assay demonstrates that TGF-b1, but not mmTGF-β induces TGFBR2 receptor internalisation. Western Blot shows whole cell extracts and surface-biotinylated fractions probed with anti-TGFBR2 and anti-E-cadherin (control) antibodies

### The presence of the co-receptor betaglycan does not impact mmTGF-β activity

Betaglycan (TGFBR3) is a co-receptor that potentiates TGF-β signalling by binding TGF-β ligands and presenting them to the signalling receptors^46^. Its role is particularly important for the TGF-β2 isoform, which has low intrinsic affinity for the signalling receptors and is largely unable to signal in the absence of betaglycan^47^. Because mmTGF-β can still bind betaglycan^46,48^, we asked whether betaglycan might influence the inhibitory activity of mmTGF-β, either by potentiating inhibition through presentation to TGFBR2, through interactions with betaglycan orphan domain, or by reducing apparent potency through ligand sequestration, through interaction with the betaglycan zona pellucida domain^46^. To investigate this, we compared wild-type NIH3T3 cells with two independent betaglycan knockout clones (BG KO1 and BG KO2), each stably expressing a CAGA_12_-mCHERRYd2 reporter. The IC₅₀ values of mmTGF-β were comparable across all tested cell lines (**Figure 5D** and **Figure S5**), indicating that the presence of betaglycan does not measurably affect the inhibitory potency of mmTGF-β. In contrast, robust potentiation of TGF-β2 signalling was observed in wild-type NIH3T3 cells relative to betaglycan-deficient clones (**Figure 5E**), consistent with the established role of betaglycan in facilitating TGF-β2 signalling.

### mmTGF-β does not affect bone morphogenic protein (BMP) signalling and does not trigger TGFBR2 internalization

TGF-β and BMP ligands are members of the same signalling protein family and engage related receptor architectures^1^. To ensure that our extensive ligand engineering did not introduce off-target interactions, we tested whether the mmTGF-β affected BMP signalling. We used an MDA-MB-231 cell line stably expressing BMP-responsive element (BRE)-luc reporter^49^. mmTGF-β showed no effect on basal reporter activity and did not attenuate BMP-4 induced signalling across a range of concentrations up to 2 µM (**Figure 5F**), whereas LDN-193189, an established kinase inhibitor of the BMP type I receptors ACVR1, BMPR1A and BMPR1B^50^, demonstrated potent inhibition at IC_50_ = 37 nM, consistent with the published literature^18^.

Ligand-induced internalization of TGF-β receptors is a well-established early event in canonical TGF-β signalling^51,52^, thus we examined whether the mmTGF-β influences receptor trafficking. Surface receptors of HaCaT cells were labelled by cell-surface biotinylation prior and after 45 min of TGF-β or mmTGF-β stimulation. TGFBR2 receptor internalization was assessed using biotin pulldown with subsequent Western blot analysis. Consistent with previous reports, stimulation with TGF-β induced internalization of TGFBR2 (**Figure 5G**). In contrast, treatment with mmTGF-β did not, indicating that receptor engagement by the inhibitor does not trigger the endocytic processes associated with pathway activation. In these assays cell surface E-cadherin was used as a control, which does not internalize upon TGF-β stimulation.

### mmTGF-β as a platform for cell-type specific inhibition

Finally, to explore whether mmTGF-β can serve as a programmable scaffold for cell-type–restricted TGF-β inhibition, we engineered a bivalent construct in which domains D4/5 of *H. polygyrus* TGF-β mimic protein 1(TGM1) (**Figure S6A**), a module known to engage CD44^53^, was connected by a flexible six residue GSGTGN linker to the C-terminus of mmTGF-β-ΔA. This design enables simultaneous targeting of TGFBR2 and CD44 at the cell surface, thereby conferring preferential activity toward cells co-expressing both receptors and an increase in potency due to avidity (**Figure 6A**). Given that CD44 is highly upregulated in activated T cells, B cells, and macrophages^54,55^, we hypothesized that dual receptor engagement would preferentially potentiate inhibition in these cell populations. To test that hypothesis, we used a model NM-18 cells that express CD44 (WT) and those knocked out for CD44 harboring the CAGA_12_-mCHERRYd2 transcriptional reporter, we observed that mmTGF-β-ΔA-TGM1-D4/5 displayed significantly enhanced inhibitory potency (ca. 30-fold) in CD44 wild-type cells compared to CD44-knockout cells. In parallel, mmTGF-β demonstrated comparable potency in both cell lines, which was also similar to the activity of mmTGF-β-ΔA-TGM1-D4/5 in CD44 knockout cells (**Figure 6B**). Western blot analysis of pSMAD2, a direct readout of pathway activity, confirmed these findings (**Figure 6B**) Together, these findings demonstrate that mmTGF-β can serve as a modular scaffold that can be functionally enhanced through fusion with auxiliary receptor-binding domains.

**Figure 6.**
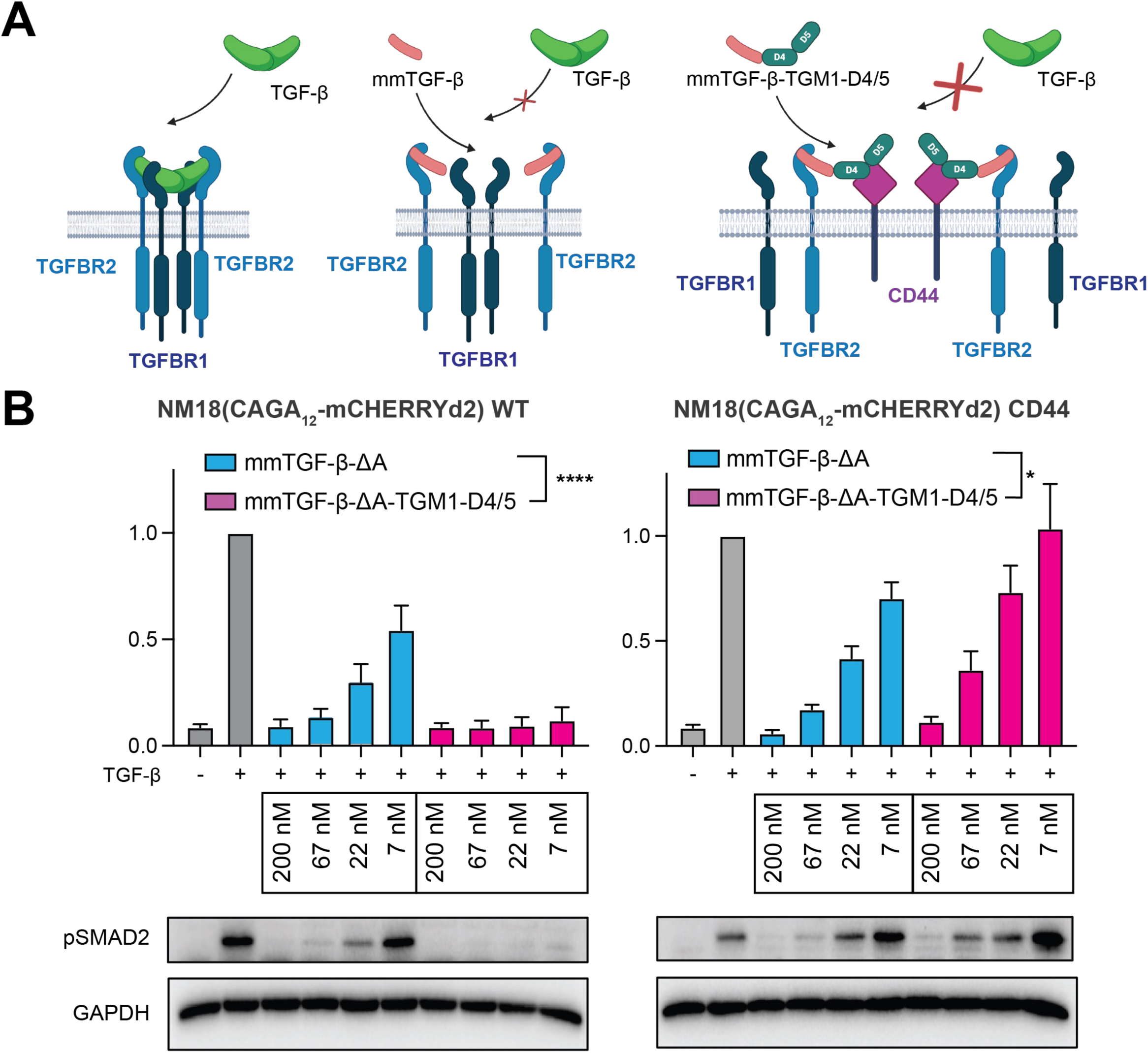
Dual engagement of TGFBR2 and CD44 confers avidity-driven inhibition of TGF-β signalling. A. Schematic illustrating the mechanism of inhibition by mmTGF-β and its bivalent variant, mmTGF-β-ΔA TGM1-D45, which simultaneously engages TGFBR2 and CD44 at the cell surface. Dual receptor targeting confers avidity-driven potency and enhances selectivity for cells co-expressing TGFBR2 and CD44. B. The potency of mmTGF-β-ΔA-TGM1-D45 was evaluated in NM18 wild-type and CD44-knockout cells harbouring the CAGA_12_-mCHERRYd2 reporter. The bivalent construct exhibited significantly enhanced inhibitory potency in wild-type cells compared with CD44-knockout (CD4 KO) cells, while showing potency comparable to mmTGF-β-ΔA in CD44-deficient cells. These results support an avidity-driven mechanism conferring CD44-dependent enhancement of inhibition. These findings were validated by Western blot analysis of phosphorylated (p)SMAD2 with GAPDH as a loading control (bottom panel). Error bars represent standard deviation. Statistical analysis was performed using GraphPad Prism (version 10.4.1). Data were analysed using two-way ANOVA.

## Discussion

The dysregulation of TGF-β signalling is well-known to promote the progression of both cancer and tissue fibrosis, yet to date no therapies targeting TGF-β have been approved for therapeutic use. There are multiple challenges to targeting the TGF-β pathway, including the secretion of TGF-β as a latent protein bound tightly to it is pro-domain, LAP^14^, storage of large quantities of TGF-β in the ECM covalently bound to fibrillin and LTBPs^27^, and the essential roles of TGF-β in tumor suppression^56^, tolerogenic immune signalling^10^, and tissue homeostasis^19^.

The majority of therapies that target TGF-β that have been explored in the clinic are systemic and target either the ligands (neutralizing antibodies or receptor traps) or receptors (kinase inhibitors and blocking antibodies)^26^. These have proven to be effective in animal models of cancer and fibrosis, yet most have failed to progress in the clinic. Through animal studies and clinical observations, one of the major liabilities that is now well-recognized is on-target toxicity, including cardiotoxicity due to interference of TGF-β mediated tissue remodeling required to sustain the heart^6,7^, as well as keratoacanthomas, due to interference of TGF-β mediated tumor suppressor activity in the skin^28^.

In recent studies, mmTGF-β, a TGF-β monomer modified to retain binding to TGFBR2, but impaired in its ability to bind and recruit TGFBR1, has been shown to potently and specifically inhibit TGF-β signalling in cultured cells^33^. This inhibitor has been further shown to have promise as an adjunct for immune therapy *in vivo*, as first shown in a carcinogen-induced model of oral cancer, where mmTGF-β delivered by osmotic pumps stimulated anti-tumor immunity and responsiveness to anti-PD-L1 checkpoint therapy^34^. In more recent studies, delivery of mmTGF-β by vaccinia, a type of OV^57^, was shown to increase regulatory regulatory Tcell fragilty and infiltration of T-cells into tumors^35^. This therapeutic approach could be further extended to potentiate responsiveness to immune checkpoint blockade in an extremely aggressive melanoma model^35^ and is now being evaluated in the clinic, both as a monotherapy using a proprietary form of vaccinia optimized to traffic to tumors^58,59^ and in combination with an α-PD-L1 checkpoint inhibitor^60^.

To optimize mmTGF-β for gene-based delivery, such as the promising OV-based therapy described above, or possibly other gene-based therapies, we sought to improve the folding of mmTGF-β when expressed in mammalian cells. To do so, we investigated simplification of the disulfide structure of mmTGF-β, as well as replacement of the cystine knot with that of PRDC, which unlike the TGF-βs, folds in a pro-domain independent manner. Though these efforts showed the accessory disulfide linking the short N-terminal helix to the β-stranded core can be removed or the cystine knot core can be swapped with that of PRDC, without impairing folding or the inhibitory potential of mmTGF-β, these modifications nonetheless led to either no or modest improvements in folding and release of inhibitory activity when mmTGF-β was expressed in mammalian cells.

Previous studies have shown that the N-terminal pro-domain of TGF-β is required to template the folding and dimerization of the C-terminal growth factor domain, and consistent with this as well as another published study which showed that dimerization of the pro-domain is not required for efficient folding, we found that a modified form of the pro-domain, lacking the bowtie knot that mediates its covalent dimerization, dramatically increased the folding efficiency. Though unprocessed pro-mmTGF-β was found to lack inhibitory activity, we found in contrast, that furin-processed pro-mmTGF-β appeared to be fully or largely unconstrained by its pro-domain, paving the way for use of furin-processible pro-mmTGF-β as an efficient means of delivering inhibitory mmTGF-β using gene-based therapies. The explanation for the full attenuation of natural TGF-β dimers by its dimeric pro-domain, but no attenuation of the inhibitory activity of mmTGF-β by its cleaved off pro-domain, likely stems from the absence of avidity in latter case, which weakens binding, and enables cell surface TGFBR2 to effectively compete mmTGF-β away from its prodomain.

Though OVs, including the engineered forms of vaccina that are being used to deliver mmTGF-β to tumors and the TME^57–59^, represent an important form of targeting that should increase the safety profile compared to a systemically-delivered TGF-β antagonist, there is also likely some targeting of bystander cells, which has the potential to lead to some on-target toxicity and thus narrow the therapeutic window. To overcome this potential barrier, we were inspired by the targeting strategy recently uncovered for the helminth parasite, *Heligmosomoides polygyrus*, which has convergently evolved modular TGF-β agonists and antagonists mimics that include, in addition to TGFBR-binding modules, additional modules that target them to co-receptors on relevant immune subsets or fibroblasts, respectively, minimizing adverse effects arising from pleiotropy^53,61–65^. To investigate the feasibility of this approach, we generated a form of mmTGF-β with the CD44 binding module of one of the parasite TGF-β mimics fused by a flexible linker to its C-terminus and found that this greatly increased the potency on target cells expressing the co-receptor, CD44. This type of approach, possibly coupled with the introduction of subtle substitutions in the TGFBR2 binding interface of mmTGF-β to attenuate the inhibitory potency of the mmTGF-β-CD44 fusion on non-target cells, therefore has the potential to increase the safety by biasing the inhibitory effect of mmTGF-β towards different classes of immune cells which are known to abundantly express the co-receptor CD44.

In previous studies, there have been efforts to attenuate TGF-β signalling on immune cell populations, for example, using bi-specific antibodies such as bintrafusp alfa, which target the immune checkpoint inhibitor α-PD-L1 on one arm and have a TGF-β receptor trap on the other arm to attenuate TGF-β signalling^66^. Though safe, this antibody failed to meet its endpoint, which was to surpass α-PD-L1 therapy alone, in a phase 3 clinical study. The underlying cause of for this failure is known, but it may be that the sequestration of TGF-β is too localized, reducing therapeutic efficacy. The choice of co-receptor targeting module to fuse to mmTGF-β must therefore be considered carefully since if the cell population being targeted is too restricted, this might also limit efficacy of the cell-targeted forms of mmTGF-β.

Overall, the data presented here collectively demonstrate that the inclusion of an engineered monomer of the pro-domain effectively templates the folding of mmTGF-β and that conjugation of mmTGF-β to co-receptor binding modules is feasible and this can confer a very strong preference for selectively inhibiting TGF-β signalling in cells harboring the co-receptor of interest. Together, these modifications are expected to further enhance the efficacy and safety of this inhibitor when delivered genetically *in vivo*.

## Materials and Methods

### Cloning

Bacterially expressed mmTGF-β variants were generated by site-directed mutagenesis of the original construct and verified by Sanger sequencing. Constructs designed for mammalian expression were synthesized and cloned into pcDNA3.1+ under the CMV promoter. All sequences were confirmed by Sanger or full-plasmid (circular consensus) sequencing. Full construct details are provided in **Table S1**.

### Cell lines

The human keratinocyte cell line (HaCaT - WT and stably transfected with GFP-SMAD2^41^), the human triple negative breast cancer (MDA-MB231) stably transfected with CAGA_12_-luciferase and BRE-luciferase reporters and the TK-Renilla reporter as an internal control^49^, human embryonic kidney cell line (HEK293T) were obtained from the Francis Crick Institute Cell Science Facility. HEK293T stably transfected with CAGA_12_-luciferase/Renilla reporter were handled at the University of Pittsburgh. Immortalized mouse embryonic fibroblast line NIH3T3 with Betaglycan KO (clone 1,2) together with WT all harboring CAGA_12_-mCHERRYd2 reporter^65^ were banked by the Francis Crick Institute Cell Services. WT and CD44 KO mouse mammary epithelial cells (NM 18) harboring CAGA_12_-mCHERRYd2^53^ were handled at the Leiden University Medical Center. All cell lines were certified negative for mycoplasma. In each case their identity was authenticated by confirming that their responses to ligands and their phenotype were consistent with published history. All mentioned cell lines were cultured in Dulbecco’s Modified Eagle Medium containing 10% FBS and were incubated at 37°C and in 5% (NM-18) or 10% CO_2_ in a humidified incubator. In addition, we used expi293 cells for p-HRV3C-mmTGF-β expression using dedicated Expi293 Pro Expression system (Gibco)

### Expression and purification of mmTGF-β variants in *E.coli*

The inhibitor variants of mmTGF-β were produced as described previously^33^. Briefly, proteins were expressed in *E. coli* BL21(DE3) in the form of insoluble inclusion bodies and after washing, the material was reconstituted in 8 M urea, reduced using DTT and refolded by dilution into non-denaturing buffer (100 mM Tris, 30 mM CHAPS, 1 M NaCl, 5 mM reduced glutathione, 20% DMSO, pH 9.5). The natively folded growth factors were isolated by high-resolution cation-exchange chromatography using a MonoS column (Cytiva 17-5166-01). Intact masses of the purified proteins were measured by LC electrospray ionization TOF MS (LC-ESI-TOF-MS, Bruker Micro TOF, Billerica, MA). mmTGF-β-PRDC and mmTGF-β-PRDCs were produced using the same strategy. However, for efficient folding a different folding buffer was required (50 mM Tris-HCl, 1 M LiCl, 500 mM Arginine, 30 mM CHAPS, 2 mM GSH and 1 mM GSSG pH 8.0).

### Expression and purification of mmTGF-β variants in HEK293T cells

The inhibitor variants of mmTGF-β and p-mmTGF-β harboring the C-terminal HA-tag were produced in HEK293T cells in DMEM + 10% FBS. 70% confluent cells were transfected with a plasmid encoding specific inhibitor variants using FuGENE HD transfection reagent. 24 h post-transfection, media was replaced with fresh DMEM + 10% FBS and expression was allowed to continue for 72 h. Conditioned media were then harvested, filtered and used for activity assays. Protein expression was assessed by Western Blott analysis using standard procedures with HRP conjugated anti-HA antibody (Roche, clone 3F10, Cat. 1203819001). The p-HRV3C-mmTGF-β variant was expressed in Expi293F cells using the ExpiFectamine^TM^ 293 Transfection Kit according to manufacturer instructions. Filtered condition media was then applied on a 5 ml cOmplete^TM^ His-Tag purification column, washed and p-HRV3C-mmTGF-β was eluted in a gradient of wash buffer containing 500 mM Imidazole. The protein was dialyzed against PBS pH 7.4, concentrated, cleaved with HRV3C protease and dialyzed against 100 mM Acetic acid. mmTGF-β was then purified on a MonoS column (Cytiva 17-5166-01) from the cleavage mixture.

### Expression and purification of TGFBR2 ECD

TGFBR2 ECD was expressed in *E.coli*, refolded and purified to homogeneity as described previously^67^.

### Expression and purification of mmTGF-β-ΔA-TGM1-CD4/5

CD44-binding domains D4/5 of TGM1 were fused onto the C-terminus of mmTGF-β-ΔA using a flexible six residue GSGTGN linker. This conjugated was produced in *E. coli* and refolded, purified to homogeneity, and native folding was validated as previously described^33^

### Luciferase reporter assays

MDA-MB-231 or HEK293T cells stably expressing CAGA_12_-Luciferase/TK Renilla or BRE-Luciferase/TK Renilla were seeded at a density of 10,000 cells/well in a 96-well plate. The following day cells were serum starved for 24 h, in the media containing 1% FBS. After starvation, cells were pre-incubated with the conditioned media/inhibitor for 30 min followed by addition of ligand (2 ng/ml TGF-β1 or 20 ng/ml BMP-4) overnight. Firefly and Renilla luciferase levels were assessed using the Dual-Luciferase reporter system according to manufacturer’s instructions and luminescence measured on the EnVision 2102 Multilabel Reader (PerkinElmer). Presented values represent Firefly/Renilla luciferase luminescence ratios, normalized to account for differences in cell number.

### Surface Plasmon Resonance (SPR) measurements

SPR binding studies were performed on a BIAcore T200 (Cytiva) and analyzed with the Biacore T200 analysis Software v3.0 at the Francis Crick Institute. TGF-β1, mmTGF-β (produced in E.coli and derived from p-mmTGF-β-HRV3C from HEK293 cells) were coupled to the surface of a CM4 chip (Cytiva) by EDC-NHS activation of the chip, followed by injection of a 1 µM solution of each ligand over the surface in sodium acetate, pH 4.5 until the RU increased by a maximum of 400 RU. All experiments were performed in 10 mM CHES, 150 mM NaCl, 3 mM EDTA, 0.01% P20 surfactant, pH 8.0 at an injection rate of 100 µl min^-1^. The higher pH was used to minimize non-specific interactions with the chip surface. The surface was regenerated in between each injection with a 10 second injection of 0.2 M guanidine hydrochloride pH 2.5. The experimental sensorgrams were obtained with double referencing with a control surface and 4 blank buffer injections. The data were analyzed by fitting the results to a 1:1 kinetic model using the SPR analysis evaluation tool (Cytiva).

### Protein folding assessment by NMR

Correct folding of mmTGF-β variants was evaluated by recording ^1^H-^15^N correlation NMR spectra and comparing their spectral fingerprints with those of the original mmTGF-β design. Protein samples were concentrated to at least 80 µM in 25 mM phosphate buffer (pH 6.0). Spectra were acquired using BEST-TROSY or HSQC pulse sequences with Watergate water suppression. All measurements were performed at 37°C on 600 or 700 MHz Bruker spectrometers equipped with TCI triple-resonance cryogenically cooled probes and running TopSpin v2.0 or v3.5. NMR data were processed using NMRPipe and analyzed with the CCPNmr Analysis software package.

### Surface biotinylation

Surface biotinylation was performed as described previously^52^. Briefly, for each condition, 2x10^6^ HaCaT cells were seeded in a 10-cm dish in DMEM supplemented with 10% FBS. After 24 h, the medium was replaced with DMEM containing 1% FBS and 24 h later, when cells had reached ∼80% confluence they were treated with either inhibitor or TGF-β for 45 min or left untreated. Following treatment, plates were transferred to ice and washed with ice-cold PBS-CM. Cells were then incubated with EZ-Link™ Sulfo-NHS-SS-Biotin (Pierce Cat. No. 21441) at 0.5 mg/ml for 45 min on ice to label surface proteins. Excess biotin was quenched by washing with 50 mM Tris-HCl pH 7.5, after which cells were harvested, lysed (1% Triton, 0.1% SDS, 10 mM EDTA, Protease inhibitors cocktail, PBS), and sonicated on ice. Lysates were cleared by centrifugation (15 min at 13 000 rpm, 4°C), and biotinylated proteins were isolated by incubation with NeutrAvidin-coated magnetic beads (Sera-Mag™ SpeedBeads, Cytiva). Captured proteins were subsequently separated by SDS-PAGE and analysed by Western Blot using standard procedures with anti-TGFBR2 (Abcam, Cat. AB259360) or anti-E-cadherin (BD Bioscience, Cat. 610181) primary antibodies, with corresponding HRP conjugated anti-rabbit (Dako, Cat. P0448) or anti-mouse (Dako, Cat P0447) secondary antibodies.

### Green Fluorescent Protein (GFP)-SMAD2 reporter assay

HaCaT cells stably expressing a GFP–SMAD2 reporter^41^ were seeded into µ-Slide 8-well chambers (ibidi) at 20,000 cells per well in DMEM supplemented with 10% FBS. After 24 h, the medium was replaced with DMEM containing 1% FBS, and cells were incubated for a further 24 h. Cells were then pre-incubated for 30 min with conditioned medium or inhibitor in the presence of the nuclear marker Hoechst 33342. The chamber was mounted on an Olympus CSU-W1 SoRa spinning-disk confocal microscope equipped with a stage-top incubator (5% CO₂, 37°C). Next, TGF-β was added to a final concentration of 2 ng/ml, and time-lapse images were acquired every 2 min for 60 min starting immediately.

Image analysis was performed using CellProfiler. Nuclei were identified in the Hoechst 33342 channel and individual cells were assigned unique IDs and tracked across all frames. Automated tracking was manually inspected and corrected when necessary. Two regions of interest were defined for each cell: (i) a nuclear region, obtained by shrinking the nuclear mask by 1 pixel to represent signal inside the nucleus, and (ii) a perinuclear cytoplasmic ring, defined as a 5-pixel-wide region starting 2 pixels outside the nuclear boundary. The nuclear–cytoplasmic ratio was calculated as the mean GFP intensity in the cytoplasmic ring divided by the mean GFP intensity in the shrunken nuclear region. Cells with a ratio greater than 1 in the first frame (indicating predominant nuclear localisation at baseline) were excluded from further analysis.

### CAGA_12_-mCHERRYd2 reporter assay (NIH3T3 cells)

WT and 2 clones of Betaglycan KO NIH3T3 cells containing CAGA-mCHERRYd2 reporter were seeded into 96-well plates (PhenoPlate^TM^-96; Revvity) at 10,000 cells per well in 150 µl DMEM supplemented with 10% FBS. After 24 h, the medium was replaced with phenol red–free DMEM containing 1% FBS, and cells were incubated for a further 24 h. Cells were then pre-treated with the indicated inhibitors for 30 min prior to TGF-β stimulation at 2 ng/ml. Fluorescence was monitored at 30-min intervals for 16 h using a Nikon Ti2 fluorescence microscope equipped with a stage-top incubator (5% CO₂, 37°C).

### CAGA-mCHERRYd2 reporter assay (NM-18 cells)

WT and CD44 KO NM-18 cells containing CAGA_12_-mCHERRYd2 reporter were seeded separately in 96-well plates (Greiner; Cat. No. 655180). The next day, cells were pre-treated with mmTGF-β or mmTGF-β-ΔA-TGM1-D45 at 37°C for 30 min prior to stimulation with 0.5 ng/ml TGF-β3 and placed in the IncuCyte S3 live-cell imaging analysis system (Sartorius). The cells were subsequently imaged every 2 h for a period of 48 h. Fluorescence intensity was analyzed using the IncuCyte software.

### Western blotting in NM-18 cells

WT and CD44 KO NM-18 cells were seeded separately in 12-well plates (Corning; Cat. No. 3512) until they reached a confluency of 80%-90% in DMEM, supplemented with 10% FBS and penicillin-streptomycin. Cells were pretreated with mmTGF-β-ΔA or mmTGF-β-ΔA-TGM1-D45 at 37°C for 30 min prior to stimulation with TGF-β3 and incubated at 37°C for 1 h. Cells were washed with phosphate-buffered saline (PBS), lysed with sample buffer, and boiled at 100 °C for 5 min. Western blotting was carried out using standard procedures. Primary antibodies used were phospho-Smad2 (Cell Signaling Technology; Cat. No. 3108) and GAPDH (Sigma-Aldrich; Cat. No. MAB374). Secondary antibodies were anti-mouse IgG (Cytiva; Cat. No. NA931) and anti-rabbit IgG (Cell Signaling Technology; Cat. No. 7074S).

### Yeast display assay

Yeast surface display clones were generated by electroporation of the *S. cerevisiae* strain EZY100 with a plasmid pCTCON2 encoding the indicated display construct. Following transformation, cells were recovered and plated on SD-CAA selective agar plates to maintain plasmid selection. A single colony of *S. cerevisiae* harbouring the indicated construct was used to inoculate 25 mL of SD-CAA medium and cultured at 30 °C with shaking overnight. Cultures reaching an OD₆₀₀ between 3 and 4 were harvested by centrifugation at 5,000 rpm for 10 min and resuspended in 25 ml of SG-CAA medium to induce protein expression. Cells were incubated for a further 16–20 h at 20 °C with shaking. On the third day, cell density was measured and 5 × 10⁷ cells were collected by centrifugation at 5,000 rpm for 10 min. Cells were washed once with 1 mL of ice-cold PBSA, and 10% of the sample was retained as an unstained control. The remaining cells were resuspended in 350 µl of ice-cold PBSA containing anti-c-Myc–FITC antibody (1:20 dilution) together with a mixture of biotinylated TGFBR2 and streptavidin–Alexa Fluor 647 and incubated for 1.5 h on ice in the dark. Following incubation, cells were pelleted, washed once with ice-cold PBSA, and resuspended in 500 µl of PBSA. Samples were transferred to FACS tubes and kept on ice until analysis. Flow cytometry data were acquired on a BD LSR II flow cytometer and analyzed using FACSDiva 8.0.1 software. All experiments were performed in three independent biological repeats.

## Acknowledgements

We thank Gerard van der Zon for technical assistance. We would like to thank Simone Kunzleman and Chloe Roustan from the Structural Biology STP at the Francis Crick Institute for assistance with SPR experiments and protein expression in expi293 cells. We are also grateful for help from the Francis Crick Institute Cell Science Facility and Advanced Microscopy Facility. This work was supported by the Francis Crick Institute (awarded to C.S.H.) which receives its core funding from Cancer Research UK (CC2021), the UK Medical Research Council (CC2021), and the Wellcome Trust (CC2021). It was also supported by a Discovery award from the Wellcome Trust (Ref. 306173 awarded to R.M.M., A.H, and P.tD), by KaliVir Immunotherapeutics (Sponsored Research Agreement awarded to A.H.), and the Dutch Research Council (NWO, Grant ID https://doi.org/10.61686/AHJBX34229 awarded to T.V. and P.tD.). This project additionally received funding from the European Union’s Horizon 2020 research and innovation programme under the Marie Skłodowska-Curie Grant Agreement No. 893196 to L.W.

---

**Figure S1.**
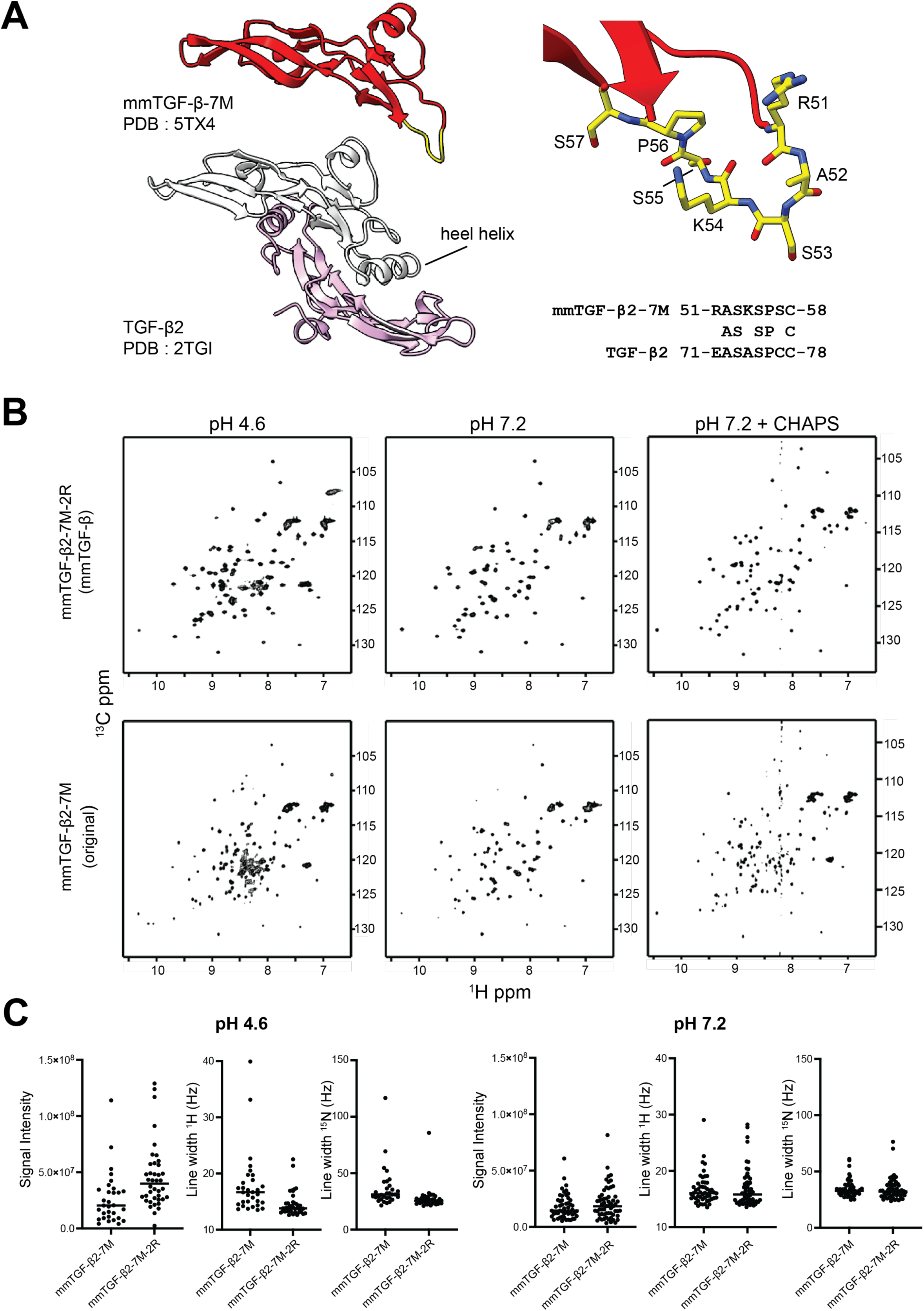
Design and initial optimization of mmTGF-β. A. Structural comparison between the active dimer of TGF-β2 and mmTGF-β2-7M. In mmTGF-β2-7M, the heel helix region is replaced by a short flexible loop (yellow), shown in detail in the zoomed-in panel on the right. B. The amide region of ^1^H-^15^N HSQC correlation spectra of the originally reported mmTGF-β2-7M and a modified construct containing two additional substitutions (S57R and S59R), termed mmTGF-β2-7M-2R and referred to in the manuscript as “mmTGF-β”. Spectra of equimolar samples were recorded at pH 4.6, pH 7.2, and pH 7.2 in the presence of CHAPS. The modified variant shows improved spectral dispersion. C. ^1^H and ^15^N signal linewidths comparison between mmTGF-β2-7M and modified mmTGF-β2-7M-2R. The modified variant shows narrower peak linewidth and stronger signals at pH 4.6 and pH 7.2 compared to the original construct, indicating improved solubility and decreased aggregation.

**Figure S2.**
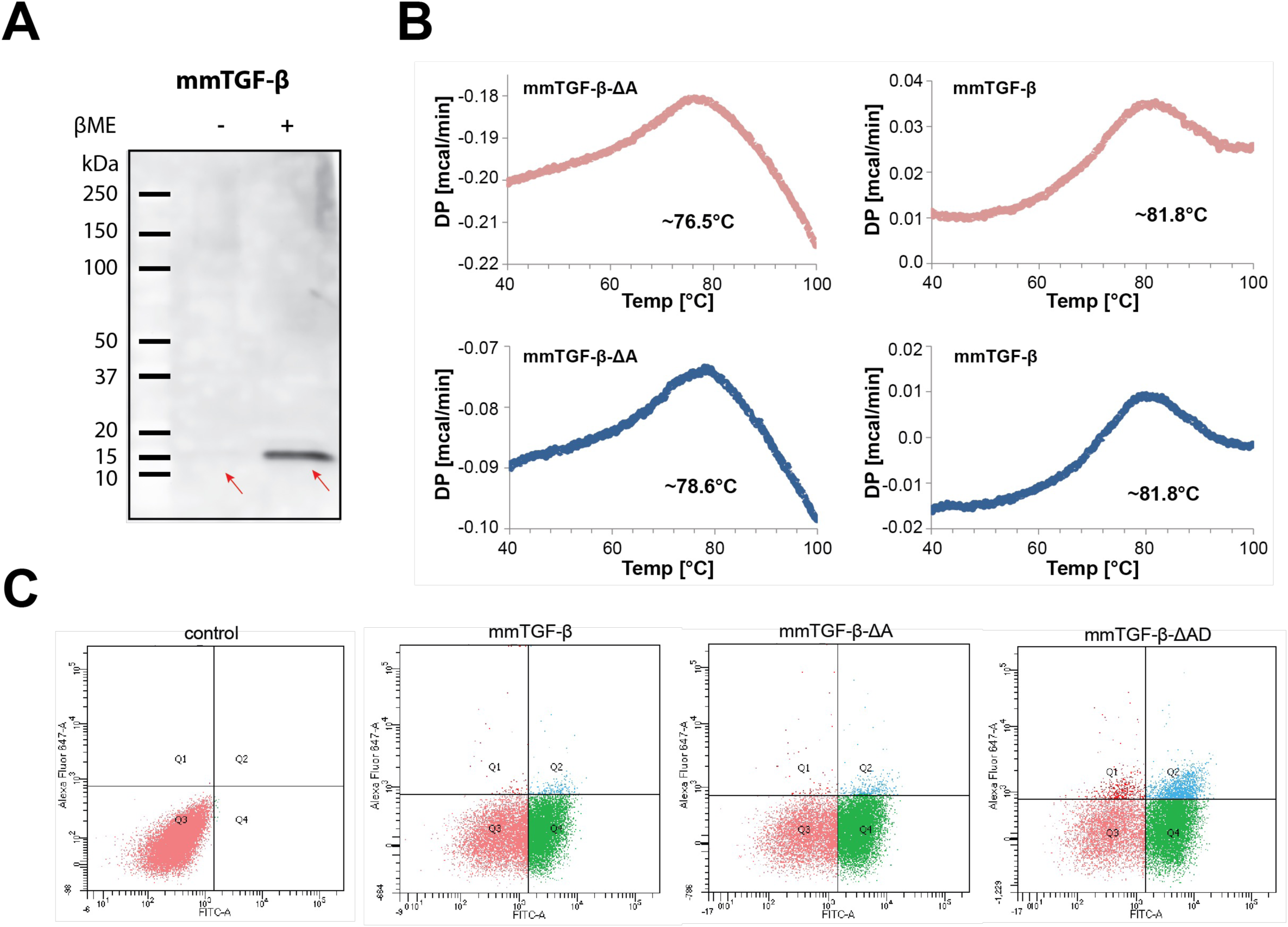
Disulfide core complexity promotes misfolding and motivates its simplification. A. Western Blot analysis of conditioned media from HEK293T cells transfected with HA-mmTGF-β construct using an antibody against the HA epitope. The band corresponding to mmTGF-β is significantly more prominent in reduced conditions, suggesting the presence of significant amounts of disulfide tethered aggregates, partially visible as a smear in the non-reducing lane. B. Differential scanning calorimetry (DSC) analysis of mmTGF-β and mmTGF-β-ΔA. Thermal unfolding profiles reveal a higher melting temperature (Tm) for mmTGF-β compared to mmTGF-β-ΔA, indicating reduced thermal stability upon ΔA substitution. Notably, both variants exhibit relatively high Tm values, consistent with preserved overall structural integrity. C. Representative flow cytometry density plots illustrating surface expression and receptor binding of yeast-displayed mmTGF-β variants. Yeast cells co-express an Aga2–mmTGF-β–GFP fusion, enabling simultaneous quantification of protein display and receptor engagement. GFP fluorescence (x-axis) reports the level of properly displayed fusion protein on the yeast surface, while AF647 fluorescence (y-axis) reflects binding of fluorescently labelled TGFBR2 extracellular domain. The unstained control defines background fluorescence in both channels. Wild-type mmTGF-β shows robust surface expression and receptor binding. In contrast, removal of a single disulfide bond modestly alters binding while largely preserving expression, whereas removal of two disulfide bonds results in enhanced receptor binding at comparable or higher levels of surface display. The double-disulfide–removed variant exhibits the strongest dual GFP and AF647 signals, indicating efficient surface presentation combined with preserved TGFBR2 engagement.

**Figure S3.**
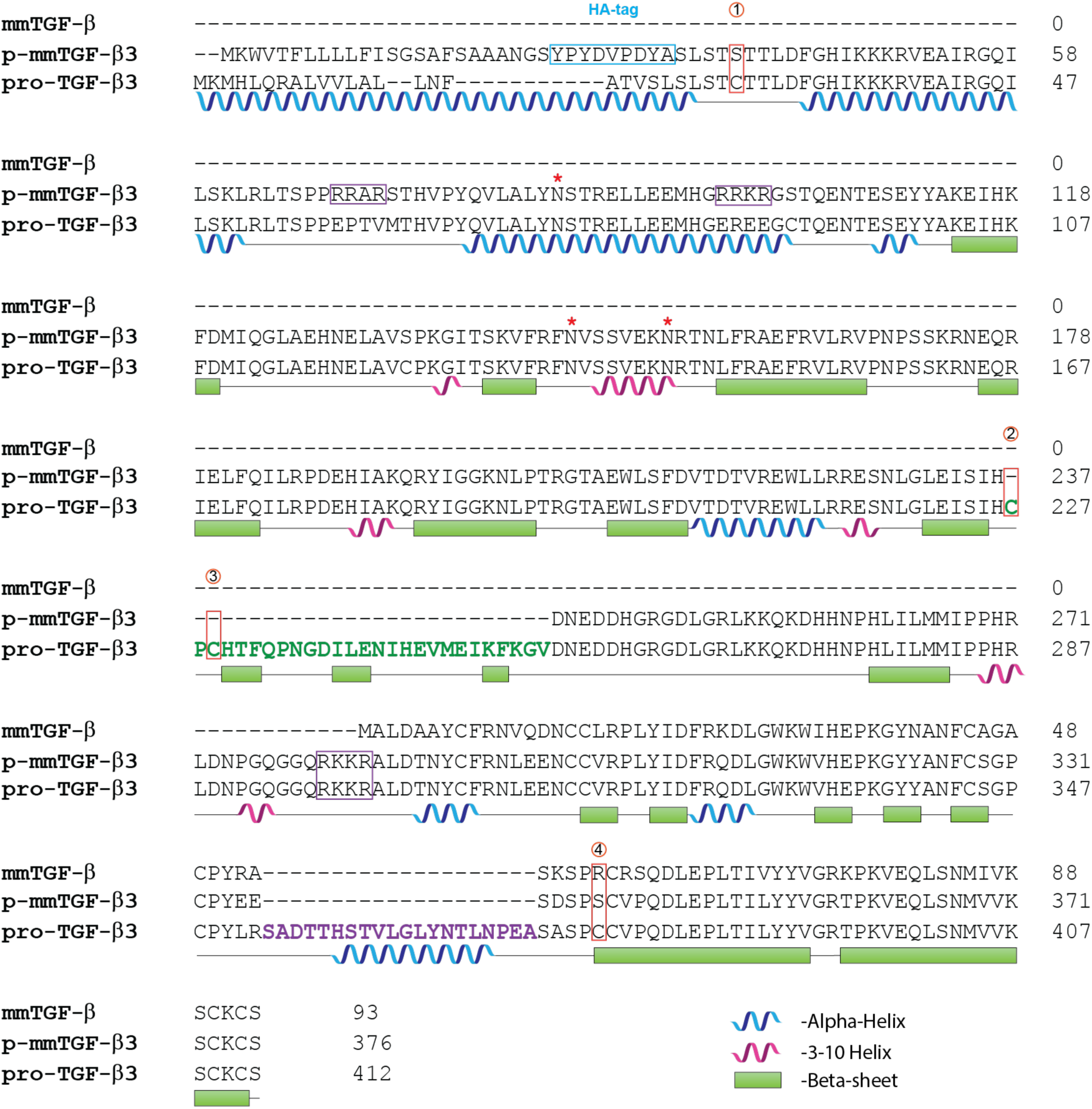
Multiple sequence alignment of wild-type pro–TGF-β3, mmTGF-β, and p-mmTGF-β3 illustrating the design logic of the engineered pro-inhibitor. Secondary structure elements are shown as blue ribbons for α-helices, red ribbons for 3₁₀-helices, and green bars for β-strands. Furin cleavage sites are highlighted with purple rectangles. The four cysteine residues removed from the growth factor and pro-domain regions are marked in red. The position of the HA tag, included to facilitate detection, is indicated with a blue rectangle. Glycosylation sites are marked by red asterisks.

**Figure S4.**
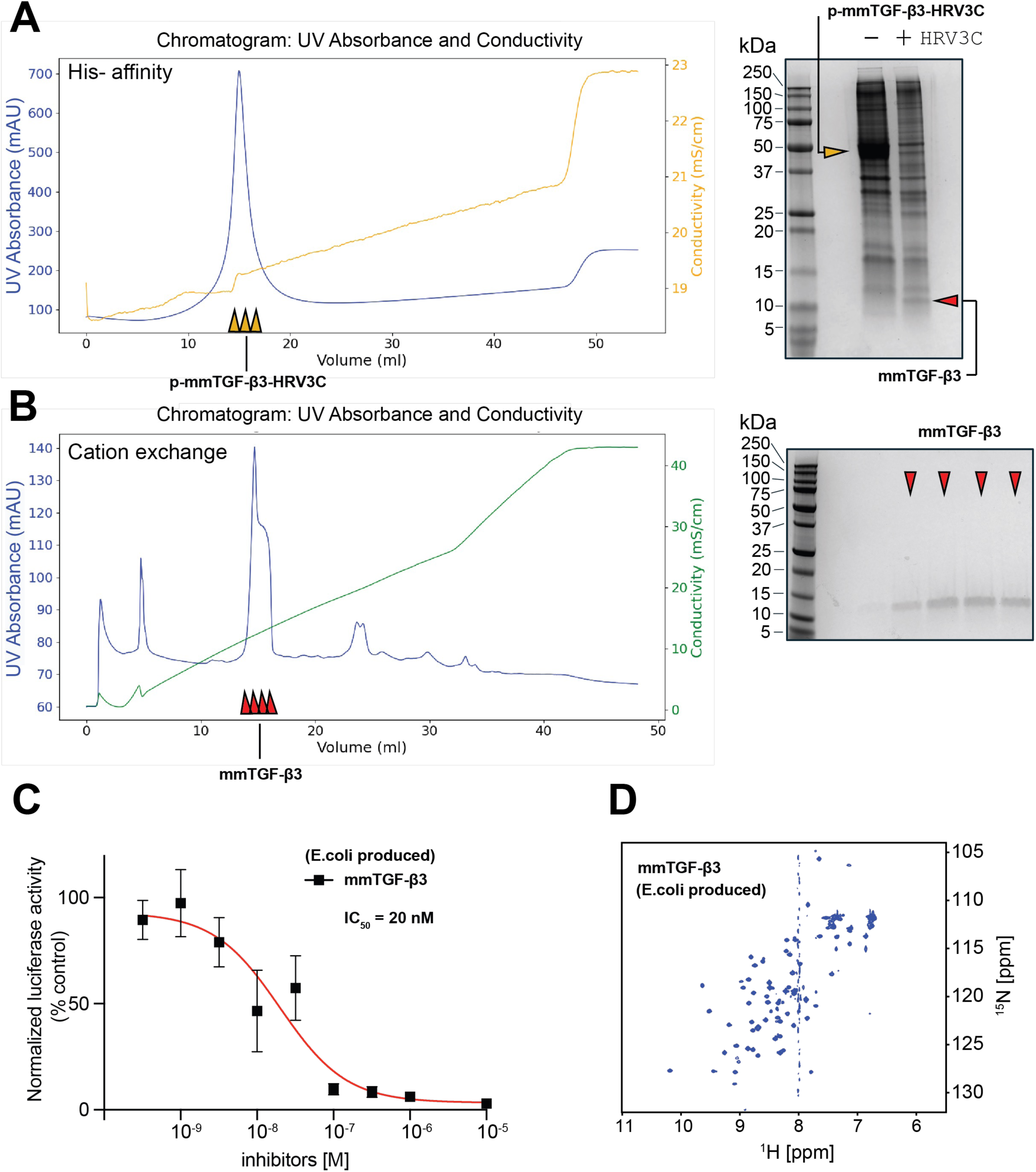
Purification of mature mmTGF-β3 from the pro-form p-mmTGF-β3. A. IMAC chromatogram of an N-terminal His-tagged p-mmTGF-β variant in which the native furin cleavage site was replaced with an HRV3C protease cleavage site. The SDS–PAGE gel on the right shows the purified uncleaved protein and the product following HRV3C digestion. B. Chromatogram from a Source S column illustrating the final purification step. Corresponding fractions containing mature mmTGF-β3 (red arrows) are shown on the elution profile and visualized on the SDS–PAGE gel to the right. C. Inhibitory activity of mmTGF-β3 (recombinantly produced in *E.coli*) measured using the CAGA_12_-luciferase/Renilla reporter assay at a constant TGF-β1 concentration and increasing inhibitor doses. D. Amide region of the ¹H-¹⁵N correlation spectra of mmTGF-β3 showing good chemical shift dispersion, consistent with a well-folded and structurally homogeneous protein. Spectra were recorded at pH 4.8 in 20 mM sodium acetate.

**Figure S5.**
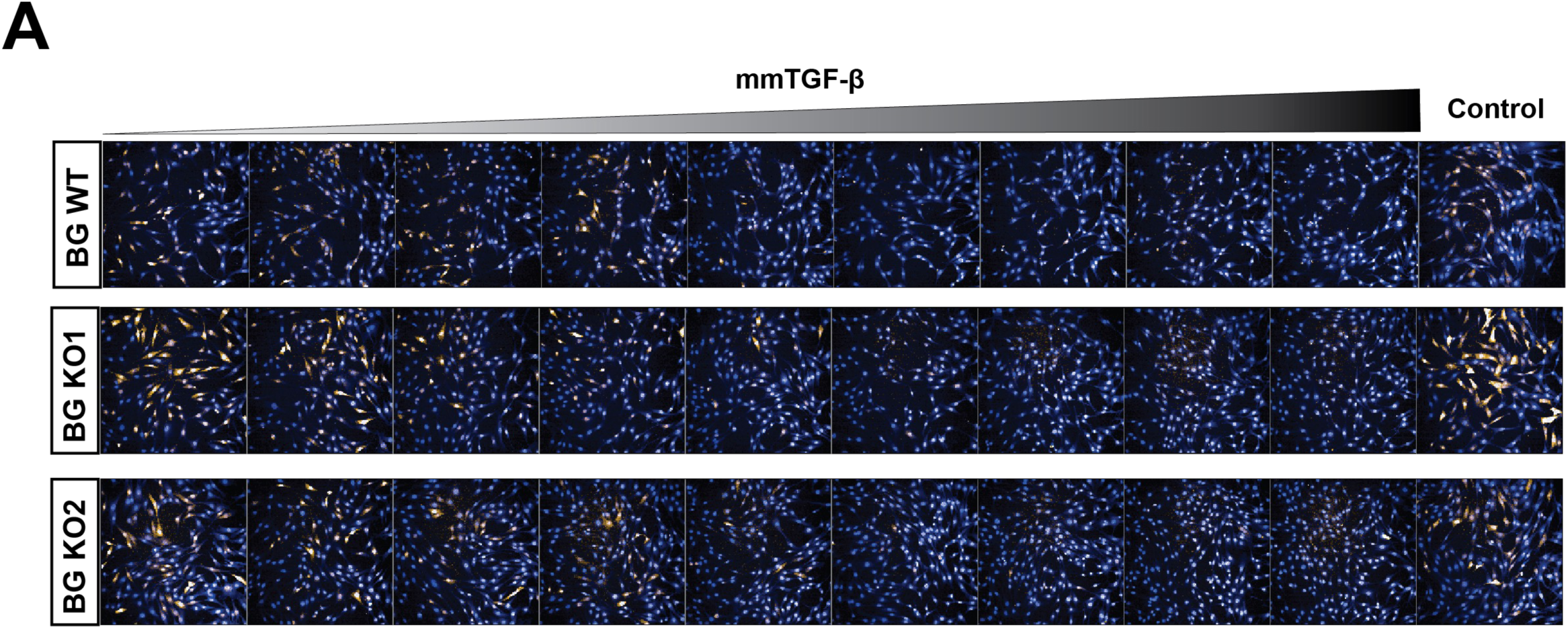
Inhibitory activity of mmTGF-β on TGF-β induced signalling in NIH3T3 wild-type and betaglycan (BG) knockout cells. Representative images show nuclei stained with DAPI (blue) and CAGA_12_-mCHERRYd2 reporter expression (orange) following TGF-β stimulation.

**Figure S6.**
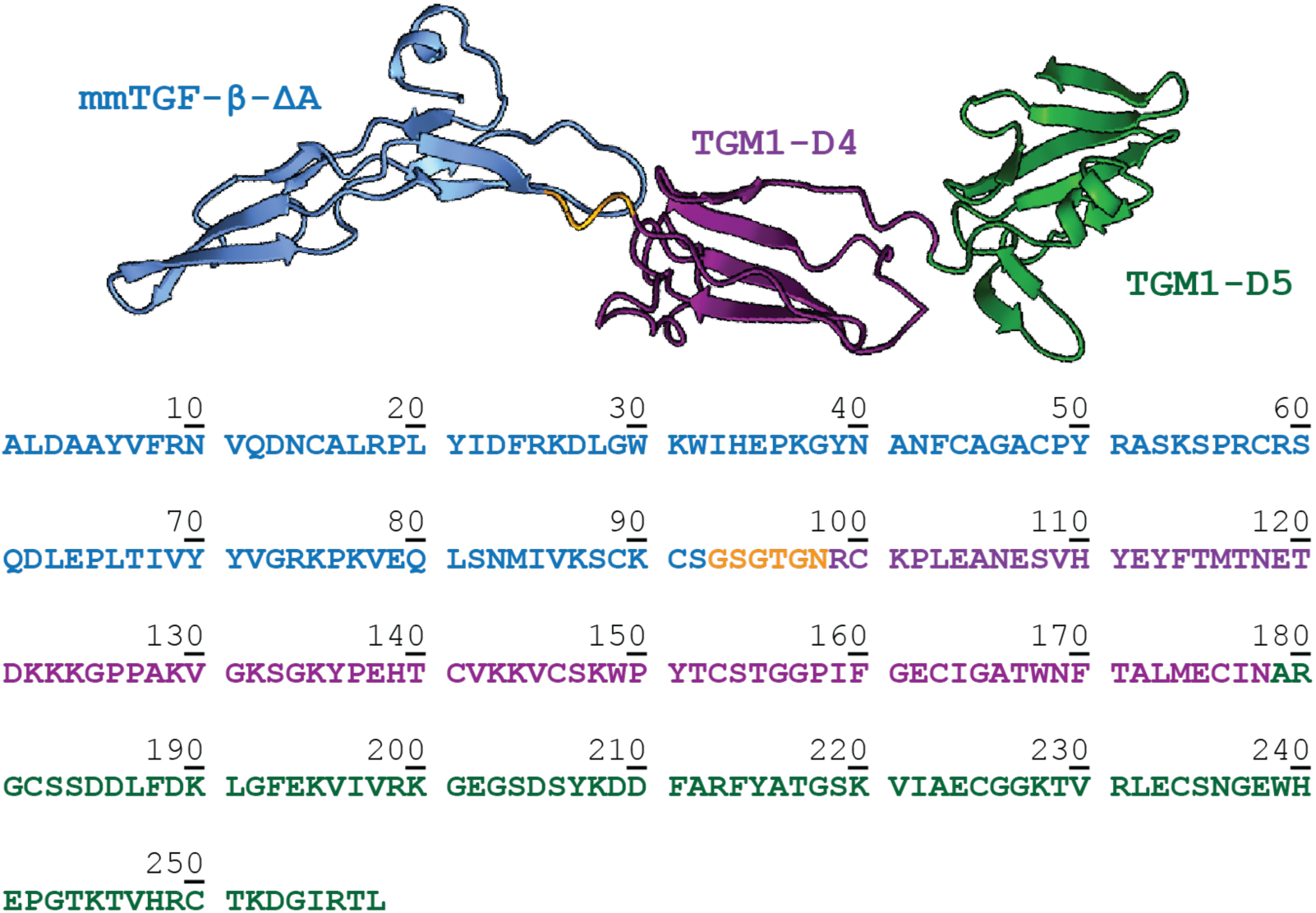
Structural model and sequence of mmTGF-β-TGM1-D4/5. Separate components are colour coded: mmTGF-β (blue), TGM1 domain D4 (purple) and TGM1 domain D5 (green), linker (orange).

**Table S1.**
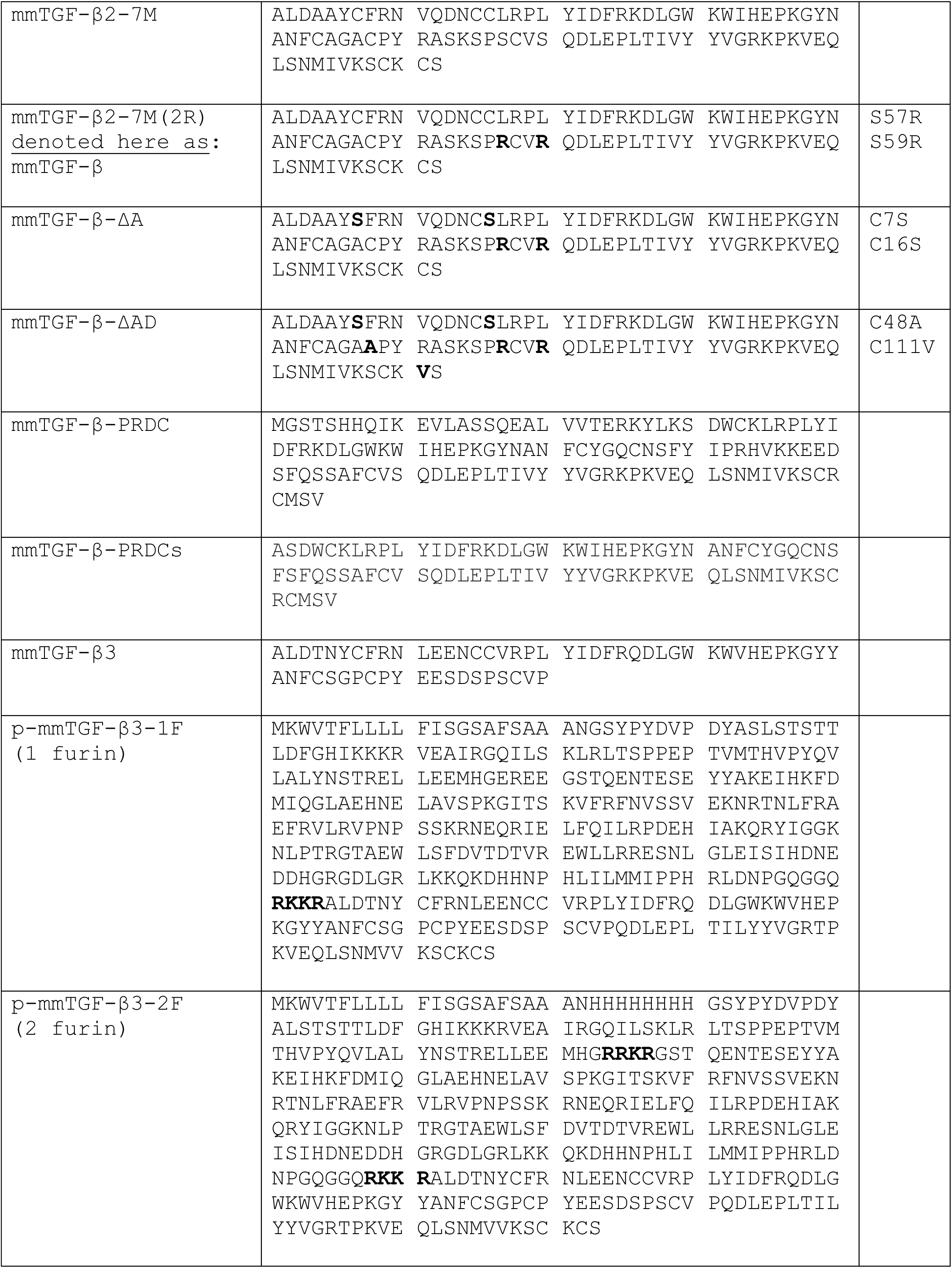
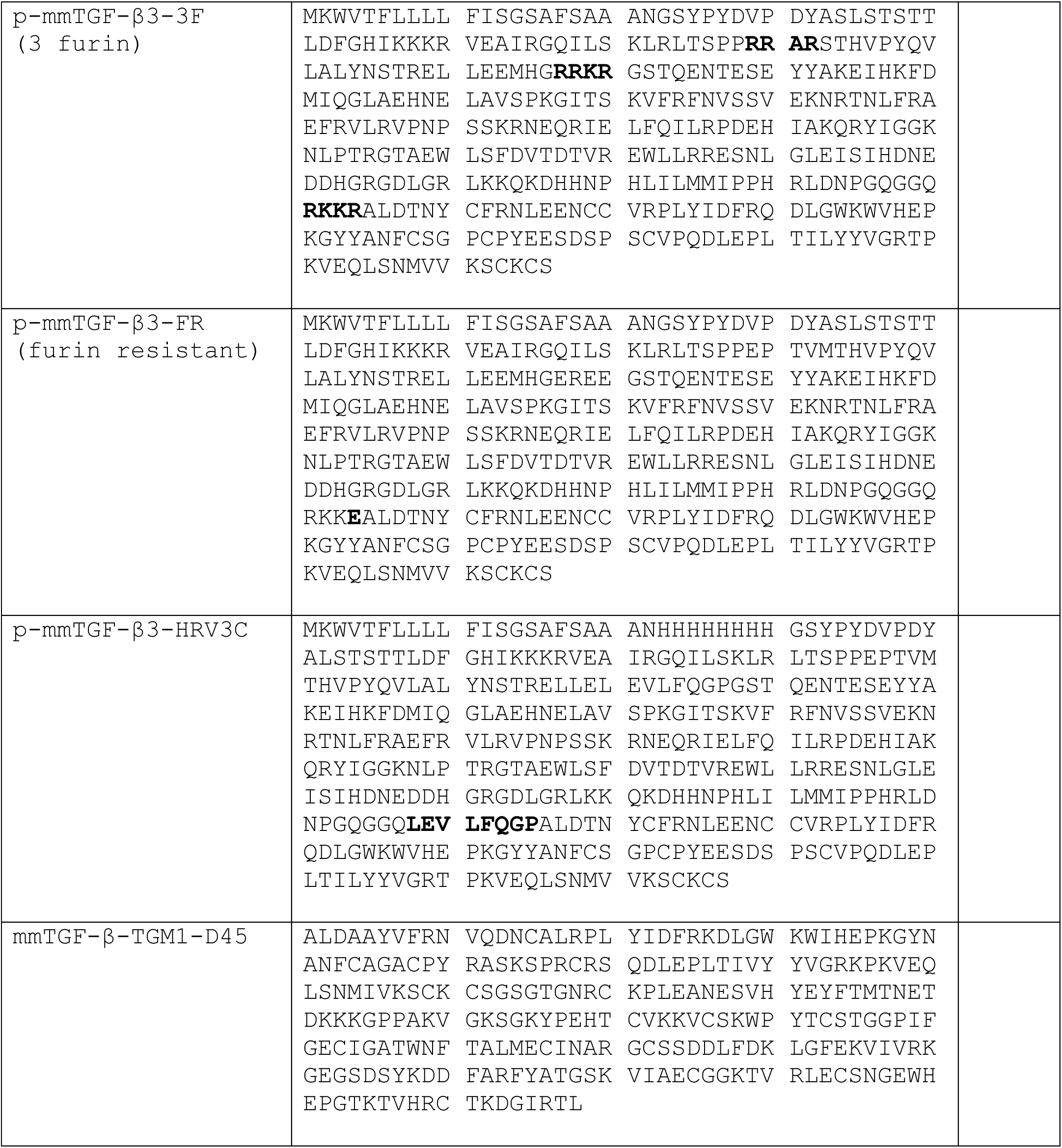
Sequences of the mmTGF-β inhibitors used in this study.

## References

1. Hinck, A.P., Mueller, T.D. & Springer, T.A. Structural Biology and Evolution of the TGF-β Family. Cold Spring Harb Perspect Biol 8, a022103 (2016).

2. Morikawa, M., Derynck, R. & Miyazono, K. TGF-β and the TGF-β Family: Context-Dependent Roles in Cell and Tissue Physiology. Cold Spring Harb Perspect Biol 8, a021873 (2016).

3. Kahata, K., Dadras, M.S. & Moustakas, A. TGF-β Family Signaling in Epithelial Differentiation and Epithelial-Mesenchymal Transition. Cold Spring Harb Perspect Biol 10, a022194 (2018).

4. Sridurongrit, S., Larsson, J., Schwartz, R., Ruiz-Lozano, P. & Kaartinen, V. Signaling via the Tgf-β type I receptor Alk5 in heart development. Dev Biol 322, 208–18 (2008).

5. Sanford, L.P. et al. TGFbeta2 knockout mice have multiple developmental defects that are non-overlapping with other TGFβ knockout phenotypes. Development 124, 2659–70 (1997).

6. Anderton, M.J. et al. Induction of heart valve lesions by small-molecule ALK5 inhibitors. Toxicol Pathol 39, 916–24 (2011).

7. Mitra, M.S. et al. A Potent Pan-TGFβ Neutralizing Monoclonal Antibody Elicits Cardiovascular Toxicity in Mice and Cynomolgus Monkeys. Toxicol Sci 175, 24–34 (2020).

8. Pardali, K. & Moustakas, A. Actions of TGF-β as tumor suppressor and pro-metastatic factor in human cancer. Biochim Biophys Acta 1775, 21–62 (2007).

9. Massague, J. TGFβ in Cancer. Cell 134, 215–30 (2008).

10. Batlle, E. & Massague, J. Transforming Growth Factor-β Signaling in Immunity and Cancer. Immunity 50, 924–940 (2019).

11. Sanjabi, S., Oh, S.A. & Li, M.O. Regulation of the Immune Response by TGF-beta: From Conception to Autoimmunity and Infection. Cold Spring Harb Perspect Biol 9, a022236 (2017).

12. Robertson, I.B. & Rifkin, D.B. Regulation of the Bioavailability of TGF-β and TGF-β-Related Proteins. Cold Spring Harb Perspect Biol 8, a021907 (2016).

13. Laiho, M., Weis, F.M., Boyd, F.T., Ignotz, R.A. & Massague, J. Responsiveness to transforming growth factor-β (TGF-beta) restored by genetic complementation between cells defective in TGF-β receptors I and II. J Biol Chem 266, 9108–12 (1991).

14. Shi, M., et al. Latent TGF-β structure and activation. Nature 474, 343–9 (2011).

15. Dong, X. et al. Force interacts with macromolecular structure in activation of TGF-beta. Nature 542, 55–59 (2017).

16. Heldin, C.H. & Moustakas, A. Signaling Receptors for TGF-β Family Members. Cold Spring Harb Perspect Biol 8, a022053 (2016).

17. Wrana, J.L., Attisano, L., Wieser, R., Ventura, F. & Massague, J. Mechanism of activation of the TGF-β receptor. Nature 370, 341–7 (1994).

18. Groppe, J. et al. Cooperative assembly of TGF-β superfamily signaling complexes is mediated by two disparate mechanisms and distinct modes of receptor binding. Mol Cell 29, 157–68 (2008).

19. Kim, K.K., Sheppard, D. & Chapman, H.A. TGF-beta1 Signaling and Tissue Fibrosis. Cold Spring Harb Perspect Biol 10, a022293 (2018).

20. Mariathasan, S. et al. TGFβ attenuates tumour response to PD-L1 blockade by contributing to exclusion of T cells. Nature 554, 544–548 (2018).

21. Tauriello, D.V.F. et al. TGFβ drives immune evasion in genetically reconstituted colon cancer metastasis. Nature 554, 538–543 (2018).

22. Sharma, P. & Allison, J.P. Immune checkpoint targeting in cancer therapy: toward combination strategies with curative potential. Cell 161, 205–14 (2015).

23. Uslu, U., Castelli, S. & June, C.H. CAR T cell combination therapies to treat cancer. Cancer Cell 42, 1319–1325 (2024).

24. Hinrichs, C.S. & Rosenberg, S.A. Exploiting the curative potential of adoptive T-cell therapy for cancer. Immunol Rev 257, 56–71 (2014).

25. Derynck, R., Turley, S.J. & Akhurst, R.J. TGFβ biology in cancer progression and immunotherapy. Nat Rev Clin Oncol 18, 9–34 (2021).

26. Akhurst, R.J. Targeting TGF-β Signaling for Therapeutic Gain. Cold Spring Harb Perspect Biol 9, a022301 (2017).

27. Robertson, I.B. & Rifkin, D.B. Unchaining the beast; insights from structural and evolutionary studies on TGFβ secretion, sequestration, and activation. Cytokine Growth Factor Rev 24, 355–72 (2013).

28. Lacouture, M.E. et al. Cutaneous keratoacanthomas/squamous cell carcinomas associated with neutralization of transforming growth factor β by the monoclonal antibody fresolimumab (GC1008). Cancer Immunol Immunother 64, 437–46 (2015).

29. Martin, C.J. et al. Selective inhibition of TGFbeta1 activation overcomes primary resistance to checkpoint blockade therapy by altering tumor immune landscape. Sci Transl Med 12, eaay8456 (2020).

30. Sawyer, J.S. et al. Synthesis and activity of new aryl- and heteroaryl-substituted 5,6-dihydro-4H-pyrrolo[1,2-b]pyrazole inhibitors of the transforming growth factor-β type I receptor kinase domain. Bioorg Med Chem Lett 14, 3581–4 (2004).

31. Richardson, L., Wilcockson, S.G., Guglielmi, L. & Hill, C.S. Context-dependent TGFβ family signalling in cell fate regulation. Nat Rev Mol Cell Biol 24, 876–894 (2023).

32. Radaev, S. et al. Ternary complex of transforming growth factor-beta1 reveals isoform-specific ligand recognition and receptor recruitment in the superfamily. J Biol Chem 285, 14806–14 (2010).

33. Kim, S.K. et al. An engineered transforming growth factor β (TGF-beta) monomer that functions as a dominant negative to block TGF-β signaling. J Biol Chem 292, 7173–7188 (2017).

34. Ludwig, N. et al. Novel TGFβ Inhibitors Ameliorate Oral Squamous Cell Carcinoma Progression and Improve the Antitumor Immune Response of Anti-PD-L1 Immunotherapy. Mol Cancer Ther 20, 1102–1111 (2021).

35. DePeaux, K. et al. An oncolytic virus-delivered TGFβ inhibitor overcomes the immunosuppressive tumor microenvironment. J Exp Med 220, e20230053 (2023).

36. Pellaud, J., Schote, U., Arvinte, T. & Seelig, J. Conformation and self-association of human recombinant transforming growth factor-beta3 in aqueous solutions. J Biol Chem 274, 7699–704 (1999).

37. Sun, P.D. & Davies, D.R. The cystine-knot growth-factor superfamily. Annu Rev Biophys Biomol Struct 24, 269–91 (1995).

38. Dennler, S. et al. Direct binding of Smad3 and Smad4 to critical TGF β-inducible elements in the promoter of human plasminogen activator inhibitor-type 1 gene. EMBO J 17, 3091–100 (1998).

39. Nolan, K. & Thompson, T.B. The DAN family: modulators of TGF-β signaling and beyond. Protein Sci 23, 999–1012 (2014).

40. Brunner, A.M., Marquardt, H., Malacko, A.R., Lioubin, M.N. & Purchio, A.F. Site-directed mutagenesis of cysteine residues in the pro region of the transforming growth factor β 1 precursor. Expression and characterization of mutant proteins. J Biol Chem 264, 13660–4 (1989).

41. Schmierer, B. & Hill, C.S. Kinetic analysis of Smad nucleocytoplasmic shuttling reveals a mechanism for transforming growth factor β-dependent nuclear accumulation of Smads. Mol Cell Biol 25, 9845–58 (2005).

42. Hata, A. & Chen, Y.G. TGF-β Signaling from Receptors to Smads. Cold Spring Harb Perspect Biol 8, a022061 (2016).

43. Jin, M. et al. Latent-TGF-β has a domain swapped architecture. Nat Commun 16, 10469 (2025).

44. Le, V.Q. et al. A specialized integrin-binding motif enables proTGF-beta2 activation by integrin alphaVbeta6 but not alphaVbeta8. Proc Natl Acad Sci U S A 120, e2304874120 (2023).

45. Jenkins, G. The role of proteases in transforming growth factor-β activation. Int J Biochem Cell Biol 40, 1068–78 (2008).

46. Wieteska, L. et al. Structures of TGF-β with betaglycan and signaling receptors reveal mechanisms of complex assembly and signaling. Nat Commun 16, 1778 (2025).

47. Cheifetz, S. et al. Distinct transforming growth factor-β (TGF-beta) receptor subsets as determinants of cellular responsiveness to three TGF-β isoforms. J Biol Chem 265, 20533–8 (1990).

48. Henen, M.A. et al. TGF-beta2 uses the concave surface of its extended finger region to bind betaglycan’s ZP domain via three residues specific to TGF-β and inhibin-alpha. J Biol Chem 294, 3065–3080 (2019).

49. Gronroos, E. et al. Transforming growth factor β inhibits bone morphogenetic protein-induced transcription through novel phosphorylated Smad1/5-Smad3 complexes. Mol Cell Biol 32, 2904–16 (2012).

50. Cuny, G.D. et al. Structure-activity relationship study of bone morphogenetic protein (BMP) signaling inhibitors. Bioorg Med Chem Lett 18, 4388–92 (2008).

51. Miller, D.S.J. et al. The Dynamics of TGF-β Signaling Are Dictated by Receptor Trafficking via the ESCRT Machinery. Cell Rep 25, 1841–1855 e5 (2018).

52. Vizan, P. et al. Controlling long-term signaling: receptor dynamics determine attenuation and refractory behavior of the TGF-β pathway. Sci Signal 6, ra106 (2013).

53. van Dinther, M. et al. CD44 acts as a coreceptor for cell-specific enhancement of signaling and regulatory T cell induction by TGM1, a parasite TGF-β mimic. Proc Natl Acad Sci U S A 120, e2302370120 (2023).

54. Baaten, B.J., Li, C.R. & Bradley, L.M. Multifaceted regulation of T cells by CD44. Commun Integr Biol 3, 508–12 (2010).

55. Zhang, Q., Wang, X., Liu, Y., Xu, H. & Ye, C. Pan-cancer and single-cell analyses identify CD44 as an immunotherapy response predictor and regulating macrophage polarization and tumor progression in colorectal cancer. Front Oncol 14, 1380821 (2024).

56. Seoane, J. & Gomis, R.R. TGF-β Family Signaling in Tumor Suppression and Cancer Progression. Cold Spring Harb Perspect Biol 9, a022277 (2017).

57. Volovat, S.R. et al. Oncolytic Virotherapy: A New Paradigm in Cancer Immunotherapy. Int J Mol Sci 25, 1180 (2024).

58. Li, F. et al. CCL5-armed oncolytic virus augments CCR5-engineered NK cell infiltration and antitumor efficiency. J Immunother Cancer 8, e000131 (2020).

59. Moon, E.K. et al. Intra-tumoral delivery of CXCL11 via a vaccinia virus, but not by modified T cells, enhances the efficacy of adoptive T cell therapy and vaccines. Oncoimmunology 7, e1395997 (2018).

60. A Study of VET3-TGI in Patients With Solid Tumors (https://ClinicalTrials.gov NCT06444815). (U.S. National Library of Medicine, 2024).

61. Johnston, C.J.C. et al. A structurally distinct TGF-β mimic from an intestinal helminth parasite potently induces regulatory T cells. Nat Commun 8, 1741 (2017).

62. Mukundan, A. et al. Convergent evolution of a parasite-encoded complement control protein-scaffold to mimic binding of mammalian TGF-β to its receptors, TbetaRI and TbetaRII. J Biol Chem 298, 101994 (2022).

63. White, S.E. et al. TGM6 is a helminth secretory product that mimics TGF-β binding to TGFBR2 to antagonize signaling in fibroblasts. Nat Commun 16, 1847 (2025).

64. Singh, S.P. et al. The TGF-β mimic TGM4 achieves cell specificity through combinatorial surface co-receptor binding. EMBO Rep 26, 218–244 (2025).

65. van Dinther, M. et al. Decoding the Mechanism of Action of a Parasite TGFβ antagonist Inspires the Creation of Cell-type-specific TGFβ Modulators. bioRxiv 10.64898/2026.02.16.706112 (2026).

66. Cho, B.C. et al. Bintrafusp Alfa Versus Pembrolizumab in Patients With Treatment-Naive, Programmed Death-Ligand 1-High Advanced NSCLC: A Randomized, Open-Label, Phase 3 Trial. J Thorac Oncol 18, 1731–1742 (2023).

67. Huang, T. & Hinck, A.P. Production, Isolation, and Structural Analysis of Ligands and Receptors of the TGF-β Superfamily. Methods Mol Biol 1344, 63–92 (2016).

